# *In situ* structure and organisation of the type IX secretion system

**DOI:** 10.1101/2020.05.13.094771

**Authors:** DG Gorasia, G Chreifi, CA Seers, CA Butler, JE Heath, MD Glew, MJ McBride, P Subramanian, A Kjær, GJ Jensen, PD Veith, EC Reynolds

## Abstract

The *Bacteroidetes* type IX secretion system (T9SS) consists of at least 19 components that translocate proteins with a type A or type B C-terminal domain (CTD) signal across the outer membrane. The overall organisation and architecture of this system including how the secretion pore (Sov) interacts with the other components is unknown. We used cryo-electron tomography to obtain the first images of the T9SS including PorK/N rings inside intact *Porphyromonas gingivalis* cells. Using proteomics, we identified a novel complex between Sov, PorV and PorA and showed that Sov interacts with the PorK/N rings via PorW and a new component PGN_1783. A separate complex comprising the outer membrane β-barrel protein PorP, PorE, and the type B CTD protein PG1035 was also identified. Similarly, the *Flavobacterium johnsoniae* PorP-like protein, SprF was found bound to the major gliding motility adhesin, SprB. Based on these data, we propose cell surface anchorage for type B CTD proteins to PorP-like proteins and a unique model where the PorK/N rings function as an outer membrane barrier to maintain the close proximity of the translocon to the shuttle and attachment complexes inside the rings, ensuring the harmonized secretion and cell surface attachment of the T9SS substrates.

## Introduction

Gram-negative bacteria have developed numerous ways to transport cargo proteins across the outer membrane (OM). To date, nine different types of secretion system (T1SS to T9SS) have been identified in bacteria ^1–3^. The type IX secretion system (T9SS), previously known as the PorSS, is specific to the Fibrobacteres-Chlorobi-Bacteroidetes (FCB) super-phylum ^4–6^. In *Flavobacterium johnsoniae*, the T9SS secretes cell-surface adhesins that are required for gliding motility ^3, 7, 8^ whilst in *Porphyromonas gingivalis*, a human oral pathogen highly associated with chronic periodontitis ^9^, the T9SS secretes cell-surface virulence factors, such as gingipains ^3^. The *P. gingivalis* T9SS secretes ∼ 30 other proteins ^5, 6, 10, 11^ including peptidylarginine deiminase (PPAD), an enzyme responsible for host protein citrullination that has been linked to rheumatoid arthritis ^12^.

Proteins secreted by the T9SS have an N-terminal signal peptide that facilitates export across the inner membrane by the Sec system and have a conserved C-terminal domain, referred to as the CTD signal, that enable them to pass through the outer membrane (OM) via the T9SS ^10, 11, 13^. Two types of CTD signal, type A and type B have been defined and these differ in the T9SS components required for their export ^14^. Most of the *P. gingivalis* secreted proteins have a type A CTD that target them to this system. Type B CTDs have recently been shown to target proteins, such as the major motility adhesin SprB, for secretion by the *F. johnsoniae* T9SS ^15^. SprB is propelled rapidly along the cell surface by the gliding motility machinery, resulting in gliding movement of the cell ^16^. Once on the surface, *P. gingivalis* type A CTD signals are removed by the sortase PorU and replaced with anionic-LPS, anchoring the proteins to the cell surface to form an electron dense surface layer (EDSL) ^17, 18^. At least some proteins with type B CTDs are also anchored to the cell surface, but the mechanism is not known.

The T9SS is comprised of at least 19 component proteins, namely PorK, PorL, PorM, PorN, Sov, PorT, PorU, PorW, PorP, PorV, PorQ, PorZ, PorE (PG1058), PorF (PG0534), PorG (PG0189), Plug (PG2092/PGN_0144), and three transcription regulators PorX, PorY and SigP ^3, 19–24^. Although the components are known, the T9SS has not been visualized *in situ*, nor has the overall organisation or architecture of this secretion system been elucidated. To date, four stable sub-complexes of the T9SS have been described: (i) the PorK-PorN-PorG complex is localised to the OM and forms a large ring-shaped structure ^25^; (ii) the PorL-PorM complex is anchored to the inner membrane ^25, 26^ with PorL extending to the cytoplasm and PorM extending to the periplasm via four domains whose structures were recently solved ^3, 27^; (iii) the attachment complex comprising PorU-PorV-PorQ-PorZ is responsible for anchoring T9SS substrates to the cell surface ^28^ and (iv) the *F. johnsoniae* translocon complexes comprising SprA (orthologue of *P. gingivalis* Sov), PorV and Plug were solved using cryo-electron microscopy ^24^. SprA forms an extremely large 36-strand transmembrane β-barrel and uses an alternating access mechanism in which the two ends of the protein-conducting channel are open at different times. SprA was found either bound laterally to the PorV OM protein or to the periplasmic Plug protein Fjoh_1759 ^24^. The spatial relationship of these four subcomplexes to each other is unknown. The remaining conserved components of the T9SS, including PorT, PorW, PorF, PorP and PorE, are poorly characterised and very little is known about their interacting partners or their roles in secretion.

In this study, we used cryo-electron tomography (cryo-ET) to visualize the T9SS in intact, frozen-hydrated *P. gingivalis* cells. We observed large rings of 50 nm radius localized to the outer membrane of the bacterium, demonstrating that the PorK-PorN rings characterised *in vitro* ^25^ also exist *in situ*. Using proteomic approaches, we then identified novel translocon complexes comprising Sov, PorV, Plug, and PorA which were linked to the PorK/N rings via PorW and PGN_1783. We further demonstrated that PorP-like proteins bind to PorE and also to type B CTD proteins including SprB which is the major motility adhesin of *F. johnsoniae*. We also identified many additional novel protein-protein interactions (PPIs) in *P. gingivalis*. This will be a useful resource to develop drug targets for this pathogen, and to explore the basic biology of *P. gingivalis* and its many relatives in the phylum *Bacteroidetes*.

## Methods

### Bacterial strains and culture conditions

*P. gingivalis* wildtype strains W50 and ATCC 33277 were grown in tryptic soy enriched Brain Heart Infusion broth (TSBHI) (25 g/L Tryptic soy, 30 g/L BHI) supplemented with 0.5 mg/mL cysteine, 5 μg/mL haemin and 5 μg/mL menadione. For blood agar plates, 5% defibrinated horse blood (Equicell, Bayles, Australia) was added to enriched trypticase soy agar. Mutant strains were grown in the same media as above with the appropriate antibiotic selections. All *P. gingivalis* strains were grown anaerobically (80% N_2_, 10% H_2_ and 10% CO_2_) at 37°C. Mutant *P. gingivalis* strains: *sov*, *porK*, *porP*, *porQ*, *porT, porW* and ABK^−^ in a ATCC 33277 background were obtained from Professor Koji Nakayama ^3^. Mutant *P. gingivalis* strains *porU*, *porV*, and *porZ* were previously produced in our laboratory in both ATCC 33277 and W50 backgrounds ^17, 29^. The mutants *porE* and W50ABK^−^*WbaP were also generated in our laboratory in the *P. gingivalis* W50 background ^18, 19^. A list of *P. gingivalis* strains used in this study is shown in **Supplementary Table 1**. *F. johnsoniae* UW101 was grown in Casitone Yeast Extract (CYE) medium ^30^ at 25^°^C with shaking. 20 µg/mL tetracycline was included in the growth medium when required.

### Sample preparation for cryo-ET

*P. gingivalis* W50ABK*WbaP ^18^ or *porK* mutant cells were harvested in the early stationary phase (OD_650_ of ∼1.0), centrifuged at 10000 g for 10 min, and resuspended in 20 mM Tris-HCl, pH 7.5 supplemented with 20 mM NaCl and 10 mM MgCl_2_. The cell suspension was then mixed with 10‐nm colloidal gold beads (Ted Pella, Inc., Redding, CA, USA) pre-coated with 5% (w/v) bovine serum albumin. A small volume (4 µL) of cells was then transferred onto freshly glow‐discharged copper R2/2 200 Quantifoil holey carbon grids (Quantifoil Micro Tools GmbH, Jena, Germany). Grids were then blotted and plunge‐frozen in a liquid ethane/propane mixture ^31^ using an FEI Vitrobot Mark IV (Thermo Fisher Scientific) and stored in liquid nitrogen.

### Cryo-ET data acquisition, processing, and subtomogram averaging

Cryo-ET data were acquired on a FEI Polara 300 keV FEG transmission electron microscope (Thermo Fisher Scientific) equipped with a Gatan energy filter and K2 Summit direct electron detector (Gatan) operating in counting mode. *P. gingivalis* ABK^−^ or PorK^−^ tilt series were acquired using UCSF tomography ^32^ or SerialEM ^33^ software packages, with a tilt range of −60° to +60°, tilt increments of 1° or 2° per tilt, a total dose of ∼155 e^−^/A^2^, and target defocus of ∼8 μm underfocus. Images were aligned, contrast transfer function corrected, and reconstructed using EMAN2 ^34, 35^. A total of 171 pre-aligned particles were submitted to PEET ^36^ for subtomogram averaging with no symmetry imposed.

### Generation of *P. gingivalis pg1035* deletion mutant and PorE-Strep

A *P. gingivalis* W50 *pg1035* mutant was produced by replacing *pg1035* codons (coding from Met to stop) with an *ermF* gene under the control of its own promoter. To do this the suicide plasmid pPG1035ermF was produced as follows. Two PCR amplicons were generated using *P. gingivalis* W50 DNA as template. PCR (1) used oligonucleotides pg1035fl_for (GGTTTGGATCCGAAGACTTC) and pg1035-ermF_rev (<u>AAGCAATAGCGGAAGCTATC</u>CCGATTGTTTTTCTTTTTTATCC) giving a 351 bp product. PCR (2) used pg1035fl_rev (GCGCTCCATACCACCACCGA) and ermF-pg1035_for (**GAAAAATTTCATCCTTCGTAG**CAAAAAACGAATCGACCTTTC) giving a 346 bp product. A third PCR (3) used the *ermF* gene in pAL30 as template with oligonucleotides ermFprom_for (<u>GATAGCTTCCGCTATTGCTT</u>) and ermFstop_rev (**CTACGAAGGATGAAATTTTTC**AGG). PCR (1) was annealed to PCR (3) by the complementary nucleotides (underlined in pg1035-ermF_rev and ermFprom_for) and spliced using pg1035fl_for and ermFstop_rev external primers producing Splice1. Splice1 was annealed to PCR (2) via complementary nucleotides (bold in ermF-pg1035_for and ermFstop_rev) and spliced using external primers pg1035fl_for and pg1035fl_rev, giving a final 1740 bp Splice2 product. Splice2 was cloned into pGem-T Easy (Promega) and integrity of the DNA confirmed by sequencing. The plasmid was linearised using SacI and transformed into electrocompetent *P. gingivalis* W50. Erythromycin resistant *P. gingivalis* recombinants were selected from agar plates supplemented with 10 µg/mL erythromycin and correct genome insertion of *ermF* at the *pg1035* locus was assessed using PCR with primers pg1035fl_for2 (agtatctcttttgtgacgag) and pg1035fl_rev2 (actcagcccttttatctcgt) that anneal external to the recombination cassette integration site.

The PorE fusion protein with a C-terminal Strep-tag™ II WSHPQFEK (PG1058StrepII) was produced for use in immunoprecipitation. To do this the suicide plasmid p1058cepA ^19^ was used as template for PCR with primers PG1058DomIIIFor1 (CAG<u>ACCCGG</u>GAATATGGGACAACCGGTC) and PG1058StrepIIRev (TTC<u>CCGCGG</u>GAGCGATTA**TTTTTCGAACTGCGGGTGGCTCCA**ACGCAACTCTTCTTC) producing an amplicon that included a 3’ portion of the pg1058 ORF with a 5’ XcmI site (underlined), and at the 3’ end, the Strep-tag™ II codons (bold), a stop codon and then a SacII site (underlined). The amplicon was digested with XcmI and SacII and ligated to XcmI/SacII-digested p1058cepA to generate p1058strepIIcepA. Plasmid p1058strepIIcepA was digested with DraIII and used to transform the *P. gingivalis* W50 *porE mutant* ^19^ with recombination at the *mfa1* locus. The *1058strepII* was under the control of the *mfa1* promoter. Transformants were selected on HBA containing erythromycin (10 µg/mL) and ampicillin (5 µg/mL). Correct genomic recombination was confirmed using PCR and one clone was designated as PorE^Strep^.

### Sov antibody

The work below was done by Genscript (www.genscript.com) according to their SC1676 package. The N-terminal (33-240) and C-terminal (2281-2499) amino acid regions of *P. gingivalis* Sov were expressed in *E. coli* and purified. Rabbit polyclonal antisera were raised against this antigen and the antigen was used to affinity purify the antibodies.

### Blue native gel electrophoresis

Blue Native gel electrophoresis was performed essentially as described ^3^. Briefly, *P. gingivalis* cells were pelleted by centrifugation at 5000 g for 5 min at 4^°^C and the pellet was suspended in native gel buffer containing 1% n-dodecyl-β-D-maltoside (DDM), complete protease inhibitors and 5 mM MgCl_2_. After sonication ^3^, the samples were clarified by centrifugation at 16,000 g for 20 min at 4^°^C. Coomassie Blue G-250 was added to the samples at a final concentration of 0.25% and the samples were electrophoresed on nondenaturing Native PAGE Novex 3-12% Bis-Tris Gels. The proteins in the gels were either transferred onto a PVDF membrane for immunodetection with antibodies specific for PorE or Sov as per the immunoblot method below ^3^ or stained with Coomassie G-250 and destained until the background was clear. For the Coomassie stained gel, the lane containing the sample was excised into 12 gel bands and in-gel digestion was performed. The gel pieces were incubated with 2% SDS, 10 mM DTT and incubated at 56^°^C for 1 h. Following incubation, 55 mM iodoacetamide was added to the gel pieces and incubated for 30 min in the dark. In-gel digestion was performed using sequencing-grade-modified trypsin (Promega) and incubated overnight at 37^°^C, as previously published ^37^. Tryptic peptides were extracted from the gel pieces using 50% acetonitrile in 0.1% TFA and sonicated for 10 min in a sonicator bath. The samples were concentrated in a vacuum centrifuge before analysis using LC-MS/MS.

### LC-MS/MS

The tryptic peptides were analysed by LC-MS/MS using the Q Exactive Plus Orbitrap mass spectrometer coupled to an ultimate 3000 UHPLC system (Thermo Fisher Scientific). Buffer A was 2% acetonitrile and 0.1% formic acid, buffer B consisted of 0.1% formic acid in acetonitrile. Sample volumes of 1 µL were loaded onto a PepMap C18 trap column (75 µM ID X 2 cm X 100 Å) and desalted at a flow rate of 2 µL/min for 15 min using buffer A. The samples were then separated through a PepMap C18 analytical column (75 µM ID X 15 cm X 100 Å) at a flow rate of 300 nL/min, with the percentage of solvent B in the mobile phase changing from 2 to 10% in 1 min, from 10 to 35% in 50 min, from 35 to 60% in 1 min and from 60-90% in 1 min. The spray voltage was set at 1.8 kV, and the temperature of the ion transfer tube was 250^°^C. The S-lens was set at 50%. The full MS scans were acquired over a m/z range of 300-2000, with a resolving power of 70000, an automatic gain control (AGC) target value of 3 × 10^6^, and a maximum injection of 30 ms. Dynamic exclusion was set at 90 s. Higher energy collisional dissociation MS/MS scans were acquired at a resolving power of 17500, AGC target value of 5 × 10^4^, maximum injection time of 120 ms, isolation window of m/z 1.4, and NCE of 25% for the top 15 most abundant ions in the MS spectra. All spectra were recorded in profile mode.

Relative abundances of proteins were quantified by MaxQuant (Ver 1.5.3.30) ^38^. Raw MS/MS files were searched against the *P. gingivalis* W50 or ATCC 33277 or the *F. johnsoniae* UW101 database with the sequence of sfGFP added. The default MaxQuant parameters were used except LFQ min ratio count was set to 1 and the match between runs was selected. MaxQuant normalised the data set as part of data processing. LFQ intensities were used for the analysis of the T9SS components and iBAQ values were used for the overall protein-protein interaction mapping in *P. gingivalis*. For quantification of the immunoprecipitated material, the LFQ intensity ratio test strain/control strain was used to identify proteins that were significantly enriched. To determine the relative abundance of the enriched proteins, the corrected iBAQ intensity (test strain-control strain) was calculated and expressed as a percentage relative to the target protein.

### Immunoblot analysis

Cell lysates from *P. gingivalis* WT lacking the gingipains (ABK^−^) and *porP* mutant were separated by SDS-PAGE and the proteins were transferred onto a nitrocellulose membrane. The membrane was blocked in 5% skim milk and probed with, mAb-1B5 (kind gift from Prof. M.A Curtis) ^39^, PorE ^19^ or Sov-specific antibodies followed by anti-mouse or anti-rabbit HRP conjugated secondary antibodies. The signal was developed using SuperSignal West Pico chemiluminescent substrate and visualised with a LAS-3000 imaging system.

### Co-immunoprecipitation

*P. gingivalis* cells were lysed in 20 mM Tris-HCl, pH 8, 100 mM NaCl, 1% DDM, Tosyl-L-Lysyl-chloromethane hydrochloride (TLCK) and complete protease inhibitors. The cells were then sonicated as per BN-PAGE protocol and clarified by centrifugation at 16,000 g for 10 min at 4^°^C. Cell lysates were mixed with either StrepTactin sepharose (GE Healthcare) for PorE^Strep^ immunoprecipation, Sov antibodies bound to agarose for Sov co-immunoprecipitation or anti-GFP agarose (Chromotek) for sfGFP_SprB (368 aa)_ immunoprecipitation. The beads were rotated on a wheel for 2 hours at 4^°^C then washed with 20 mM Tris-HCl, pH 8, 100 mM NaCl, followed by high salt washes and lastly with 10 mM Tris-HCl, pH 8. The beads were suspended in Laemmli buffer and boiled for 10 min. The samples were subjected to SDS-PAGE electrophoresis for a short period of time and stained with Coomassie Blue. The sample was excised as a single band and subjected to tryptic digestion and LC-MS/MS analysis.

## Results

### Cryo-electron tomography

To study the structure of the T9SS *in vivo*, we imaged *P. gingivalis* cells using cryo-ET. We imaged a *P. gingivalis* strain that lacks gingipains and A-LPS (W50ABK*WbaP), but has a fully functional T9SS ^18^, because the absence of the electron-dense surface layer (EDSL) created by the attachment of gingipains and other CTD proteins to the cell surface via A-LPS 18 makes identification easier in cellular cryotomograms (Figure 1A). In our cryotomograms of the W50ABK*WbaP strain, we observed pairs of densities in the periplasmic space associated with the outer membrane that were roughly 50 nm apart (Figure 1C, D, and E), consistent with the diameter of purified PorK-PorN rings previously obtained by single particle reconstruction methods ^25^. Rotating by 90° about an axis in the plane of the membrane produces top views. In this orientation, the two densities previously seen in the side-view appear as two disconnected arches (Figure 1F, G, and H). This is consistent with a ring-like structure affected by the tomographic missing wedge, caused by a limitation in the angular range accessible during tilt-series acquisition ^40^. These rings were always found in direct contact with the outer membrane, but at varying distances away from the inner membranes, suggesting that the assembled state of the PorK-PorN moiety does not depend on interactions with inner membrane-bound components. Particles were found in 91% of the 80 cells imaged. While no particles were found in 7 cells, 6 particles were found in a single cell. The average number of particles found per cell was 2.1 (Figure 1B).

**Figure 1:**
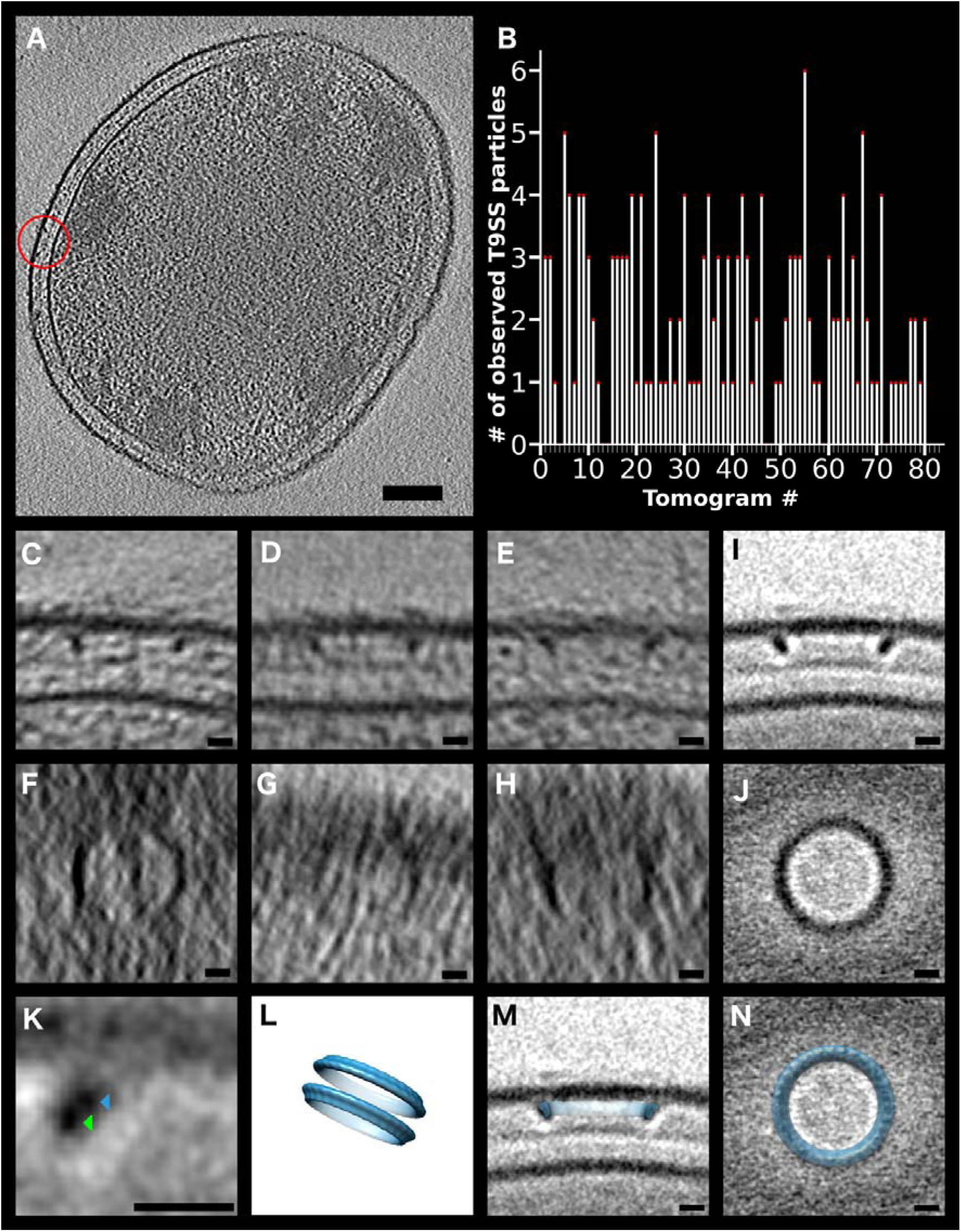
*In situ* images of the *P. gingivalis* type IX secretion system PorK-PorN. A. Slice through a cryotomogram of an intact frozen-hydrated *P. gingivalis* W50ABK*WbaP cell lacking the EDSL. A T9SS particle is circled in red. Scale bar: 100 nm. B. Number of observed T9SS particles per tomogram. C, D, E: Tomographic slices through 3 different T9SS particles in a side-view orientation (scale bar: 10 nm) F, G, H: Tomographic slices through 3 different T9SS particles in a top-view orientation (scale bar: 10 nm) I: Subtomogram average from 171 particles (side-view). Scale bar: 10 nm. J: Subtomogram average from 171 particles (top-view). Scale bar: 10 nm. K: Enlarged tomographic slice through the subtomogram average showing one side of ring (side-view). Two densities are seen: PorK (blue arrow) and PorN (green arrow). Scale bar: 10 nm. L: Previous single particle reconstruction of 107 purified PorK-PorN particles (from reference 25). M, N: PorK-PorN *in vitro* ring single particle reconstruction superimposed on M: side view or N: top view of the *in situ* subtomogram average.

No particles were found in 23 tilt-series of the *porK* mutant lacking PorK, the outer-membrane-bound lipoprotein. The fact that no rings were found in tomograms of the *porK* mutant, combined with the essentially perfect match between the *in situ* densities and the *in vitro* reconstruction of purified PorK-PorN rings ^25^, as well as the correct localization to the outer membrane ^3^, strongly indicate that these rings are the PorK-PorN components of the T9SS.

The initial sub-tomogram average did not exhibit a completely circular ring. In order to improve the resolution, we focused our alignment to just the region near the outer membrane and increased the number of particles to 171. In this improved average (Figure 1I and J) two densities were resolved within the ring (Figure 1K). The larger ring lay adjacent to the outer membrane and exhibited an outer and inner diameter of 50 and 37 nm, respectively. The smaller ring was found beneath the larger ring and had an outer and inner diameter of 45 and 35 nm, respectively. Similar substructures were seen previously in the single particle reconstruction of purified PorK-PorN rings (Figure 1L) ^25^. Unfortunately we could not resolve individual monomers within the rings, so we could not discern the symmetry. The previously published reconstruction of PorK-PorN rings superimposes well on the side and top views of the *in situ* subtomogram average (Figure 1M and N). No other components of the T9SS were clearly resolved, presumably because of either structural or compositional heterogeneity.

### The *P. gingivalis* protein-protein interactome

Having been unable to resolve other components of the T9SS in our subtomogram average, we investigated the global protein-protein interaction (PPI) network of the *P. gingivalis* ABK^−^ strain using Blue-Native gel electrophoresis (BN-PAGE)/mass spectrometry. Analysis of PPI networks is a powerful approach to dissect protein function, potential signal transduction, and virulence pathways. We selected the ABK^−^ strain for the PPI study for several reasons; firstly, the three gingipains are amongst the most abundant proteins in *P. gingivalis* and hence their elimination allowed us to detect many less abundant proteins by BN-PAGE. Secondly, the gingipains are proteases and their removal minimized protein degradation in the cell lysate. Thirdly, the gingipains process some of the *P. gingivalis* surface proteins, affecting the protein profile of the cell. Finally, since T9SS mutants lack active gingipains ^3, 17, 29^, a *P. gingivalis* strain that does not produce gingipains but has a functional T9SS is useful to compare with the T9SS mutants in studies of this secretion system. BN-PAGE was performed with *P. gingivalis* cells (four biological replicates) lysed in 1% DDM. After electrophoresis each lane was excised into 12 gel bands (Figure 2A) and subjected to tryptic digestion, mass spectrometry and MaxQuant analysis. A total of 1232 proteins were identified and the protein abundances (iBAQ) were plotted as a heat map for each gel band (Figure 2B, Supplementary Table 2). Proteins with a similar native migration profile may be part of the same protein complex. Using this analysis, a large number of potential PPIs were seen.

**Figure 2:**
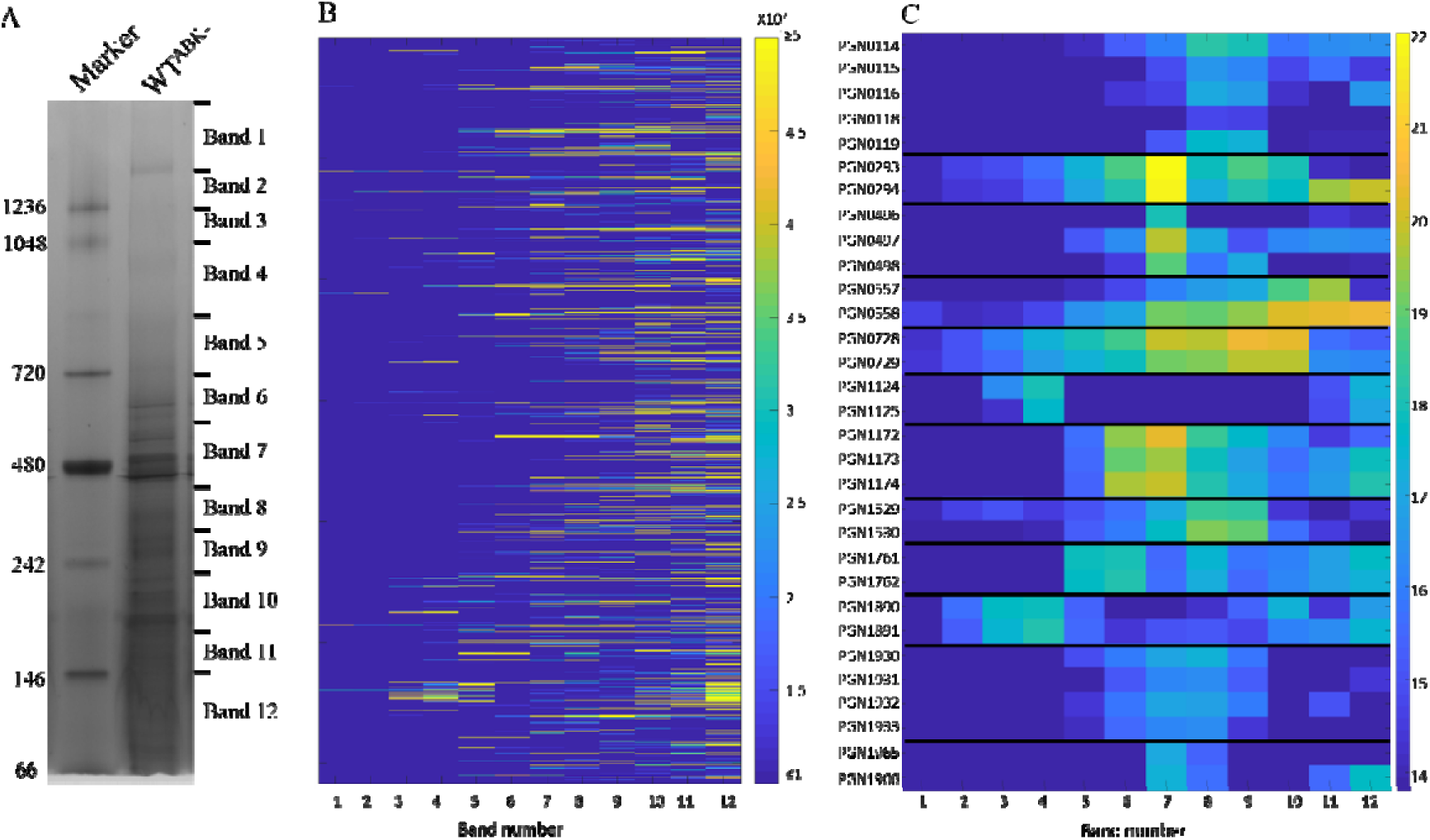
Native migration profiles of *P. gingivalis* proteins plotted as a heat map. (A) *P. gingivalis* cells lacking the gingipains (ABK^−^) were lysed in 1% DDM, electrophoresed on a BN-PAGE gel and stained with Coomassie blue. The gel lane was sliced into 12 bands and an in-gel tryptic digestion was performed on the gel pieces. Tryptic fragments were analysed by mass spectrometry and identified using MaxQuant software. (B) Migration profile overview of all the proteins identified in each band of the BN-PAGE gel based on the iBAQ values (average of four replicates). (C) Migration profiles of selected complexes in each band with the iBAQ values expressed on a natural log scale. See also **Supplementary Table 2**.

Twelve potential protein complexes consisting of proteins expressed from adjacent genes are shown in Figure 2C and described below. The putative components of a *P. gingivalis* 11 respiratory chain were predicted from bioinformatic studies, and proposed to consist of six subunits (PG2177/PGN_0119-PG2182/PGN_0114) ^41^. Previously, two of these, PGN_0119 and PGN_0116 were shown to interact ^42^. Here we observed all but one of the subunits to peak at bands 8 and 9 suggesting at least five are part of one complex. Similarly, RagA and RagB (PGN_0293, PGN_0294) co-migrated by BN-PAGE, as did Omp40 and Omp41 (PGN_0728, PGN_0729) (Figure 2C). Direct interactions between these protein pairs were reported previously ^43–45^. Several other sets of proteins also had migration profiles that suggested they formed complexes: PGN_1124-PGN_1125, PGN_1172-PGN_1174, PGN_1529-PGN_1530, PGN_1890-PGN_1891, PGN_1930-PGN_1933 and PGN_1965-PGN_1966 (Figure 2C). Interactions between the proteins in these groups have not been reported previously, but putative homologues for some of these proteins are thought to interact in other species according to STRING ^46^. These potential PPIs are a useful resource for studies of *P. gingivalis* physiology.

### Blue-native PAGE proteomic analysis of *P. gingivalis* (ABK^−^) identifies co-migrating proteins of the T9SS

Using the data above we traced the native profiles of the T9SS proteins to gain insights on the T9SS interactome. The Label Free Quantification (LFQ) intensities of the T9SS proteins were normalised and plotted (Figure 3A-D). PorK and PorN were the predominant T9SS proteins observed in gel bands 1, 2 and 3. In gel bands 4 and 5, PorK and PorN intensities were increased and joined by PorL and PorM (Figure 3A). PorT was found in gel bands 1 to 5 following a similar profile to PorK and PorN suggesting it may interact with the PorK-PorN complex (Figure 3A). However, the majority of PorT was found in Band 12. PorG has previously been shown to interact with PorK and PorN ^25^. We were unable to identify PorG in ABK^−^, but PorG had a similar migration profile as PorK and PorN in the *porV*, *porU, porP* and *porE* mutants (**Supplementary table 3**). In addition, the following groups of proteins had BN-PAGE migration profiles that suggested possible protein complexes: (i) Sov, PorW and a putative T9SS component PGN_1783 (Figure 3B), (ii) PorP, PorE, and PorF (Figure 3C) and (iii) the attachment complex comprised of PorU, PorV, PorQ and PorZ (Figure 3D). These are discussed in greater detail below.

**Figure 3:**
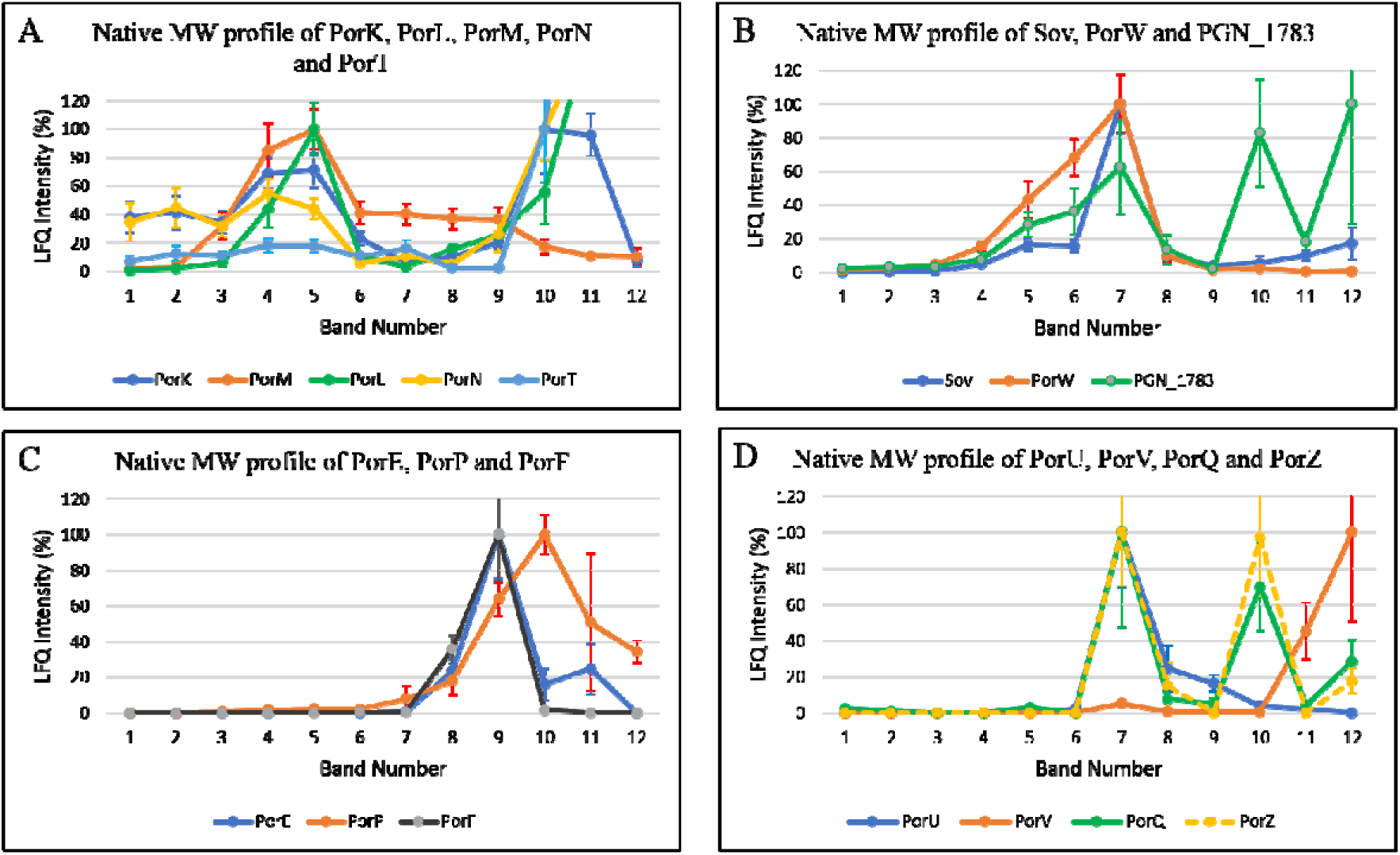
Putative sub-complexes of the T9SS identified by Blue-Native PAGE analysis of *P. gingivalis* ABK^−^. The LFQ intensities for the T9SS proteins identified from each BN-PAGE band were converted to percentage with the highest LFQ intensity being assigned 100% for each protein in the group except for PorT, PorL and PorN where the intensity at band 10 was assigned 100% to aid visualisation of the high molecular weight bands. Proteins that co-migrated on a BN-PAGE gel were grouped and plotted together (A) PorK, PorL, PorM, PorN and PorT, (B) Sov, PorW and PGN_1783 (C) PorE, PorP and PorF, (D) PorU, PorV, PorQ and PorZ

### The PorQ-PorZ sub-complex is formed independently of PorU and PorV

A peak of PorU, PorV, PorQ and PorZ was observed in gel band 7 in the BN-PAGE analysis of *P. gingivalis* ABK^−^ (Figure 3D). This suggests an interaction between PorU-PorV-PorQ-PorZ consistent with our previous findings where we named it the attachment complex ^28^. We also observed PorQ and PorZ peaks at gel band 10 (Figure 3D) suggesting the presence of a PorQ-PorZ sub-complex. To investigate which T9SS components are essential for the formation of the attachment complex or the PorQ-PorZ sub-complex, we performed BN-PAGE analysis on *P. gingivalis* T9SS mutants (Figure 4). The attachment complex was not observed in any of the mutants, but the putative PorQ-PorZ sub-complex was observed in *porU* and *porV* mutants suggesting that the PorQ-PorZ complex can form in the absence of PorU and PorV.

**Figure 4:**
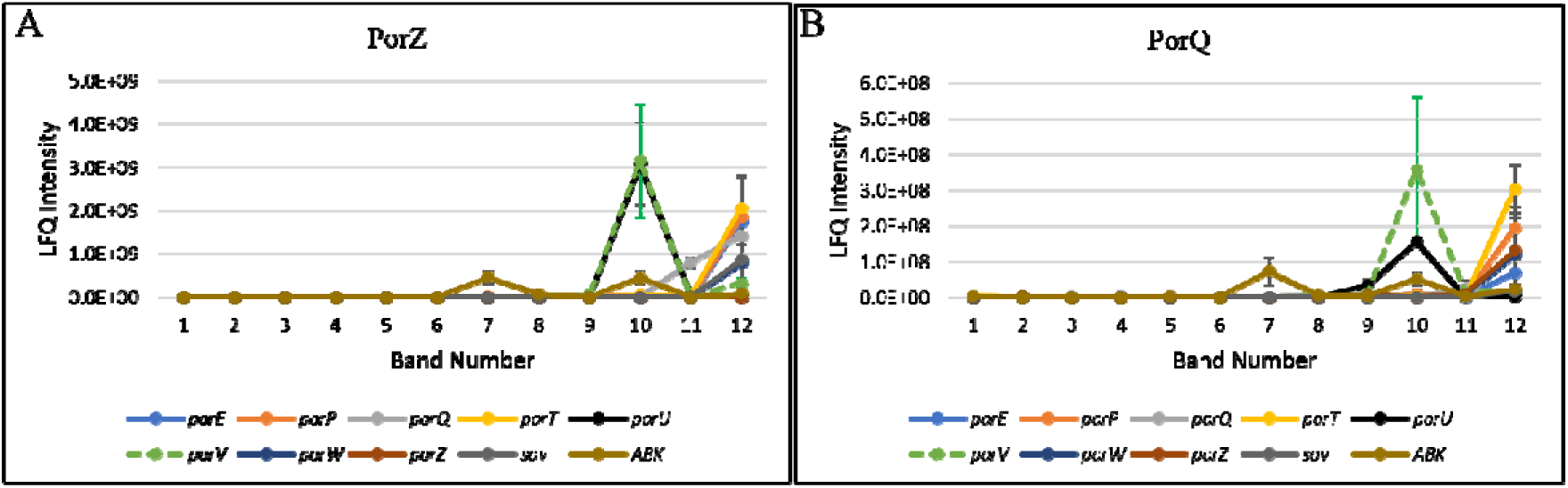
The PorZ and PorQ complex is formed independently of PorU and PorV. *P. gingivalis* T9SS mutants (*porT*, *porQ*, *porP*, *porE*, *porW*, *sov*, *porZ*, *porU* and *porV*) were lysed in 1% DDM and electrophoresed on a 3-12% BN-PAGE gel. The gel lanes were sliced at the same positions as in Fig 2A. The tryptic fragments were analysed by mass spectrometry and MaxQuant software. The LFQ intensities of (A) PorZ and (B) PorQ in each T9SS mutant and ABK^−^ were plotted.

### High molecular weight complexes of PorV observed in *porU*, *porQ* and *porZ* mutants

Previously, an accumulation of PorV bound to T9SS substrates (CTD proteins) was observed in the *porU* mutant ^28^. In the BN-PAGE analysis of strain ABK^−^, a minor PorV peak was observed at gel band 7, where it is assumed to be part of the attachment complex (Figure 3D). However, the majority of PorV was found in gel band 12, which is likely to be the free form of PorV that is not bound to any other protein (Figure 3D). In contrast, in the *porU*, *porQ* and *porZ* mutants larger PorV complexes predominated (Table 1). These high molecular weight complexes may contain PorV and CTD proteins stalled in the cleavage and surface attachment process. PorU is required for CTD signal cleavage and attachment ^17, 18^. The results above are consistent with this and suggest that PorQ and PorZ are also required for the surface attachment of CTD proteins.

**Table 1:** The LFQ intensity values of PorV protein in the T9SS mutants show high molecular weight complexes of PorV in *porU*, *porQ* and *porZ* mutants. The highest LFQ intensity in each mutant has been assigned 100%.

|  | ABK<sup>-</sup> | *porW* | *sov* | *porT* | *porP* | *porE* | *porZ* | *porU* | *porQ* |
| --- | --- | --- | --- | --- | --- | --- | --- | --- | --- |
| **Band 1** | 0 | 0 | 0 | 0 | 0 | 0 | 0 | 1 | 0 |
| **Band 2** | 0 | 0 | 0 | 0 | 0 | 0 | 0 | 0 | 0 |
| **Band 3** | 0 | 0 | 0 | 0 | 0 | 0 | 0 | 0 | 0 |
| **Band 4** | 0 | 0 | 0 | 0 | 0 | 0 | 0 | 0 | 0 |
| **Band 5** | 0 | 0 | 0 | 0 | 0 | 0 | 1 | 3 | 1 |
| **Band 6** | 0 | 0 | 0 | 0 | 0 | 0 | 3 | 4 | 32 |
| **Band 7** | 3 | 2 | 0 | 0 | 2 | 0 | 18 | 23 | 75 |
| **Band 8** | 0 | 1 | 1 | 1 | 1 | 0 | 30 | 25 | 33 |
| **Band 9** | 1 | 0 | 0 | 0 | 0 | 0 | 41 | 12 | 77 |
| **Band 10** | 0 | 0 | 0 | 0 | 1 | 0 | 22 | 95 | 26 |
| **Band 11** | 11 | 13 | 19 | 10 | 4 | 6 | 100 | 100 | 100 |
| **Band 12** | 100 | 100 | 100 | 100 | 100 | 100 | 74 | 3 | 1 |

### PorW is required for Sov to form the 750 kDa complex

The Sov, PorW and PGN_1783 proteins had similar native profiles showing increased amounts between gel bands 5 to 7, suggesting that these proteins may interact to form a complex (Figure 3B). To further explore the potential interactions of these three proteins, we examined the BN-PAGE data of the T9SS mutants. The small Sov peak at gel band 5 disappeared in the *porW* mutant while the peak at gel band 7 remained suggesting that gel band 5 may include a complex containing Sov and PorW (Figure 5A). The PorW peak was not apparent in the *sov* mutant (Figure 5B) and the PGN_1783 peak disappeared in both *porW* and *sov* mutants (Figure 5C) consistent with all three proteins having some dependence on each other. While Sov, PorW, and PGN_1783 peaked at gel band 7 in the ABK^−^ strain, in some mutants they peaked at band 5 (**Supplementary Figure 1**). It was observed that all mutant strains exhibiting a prominent Sov peak at band 5 (*porT, porP, porE*) also exhibited prominent peaks of PorW and PGN_1783 at band 5, while the remaining mutants (*porU, porV, porZ*) which did not produce a prominent Sov peak at band 5 also did not produce prominent peaks of PorW and PGN_1783 at band 5 (**Supplementary Figure 1**). The close correlation of these proteins supports the presence of a Sov-PorW-PGN_1783 complex in band 5. The peak at band 7 may include a PorW-PGN_1783 complex, but the Sov in band 7 appears to be an independent complex, because it was also present in the *porW* mutant (Figure 5A).

**Figure 5:**
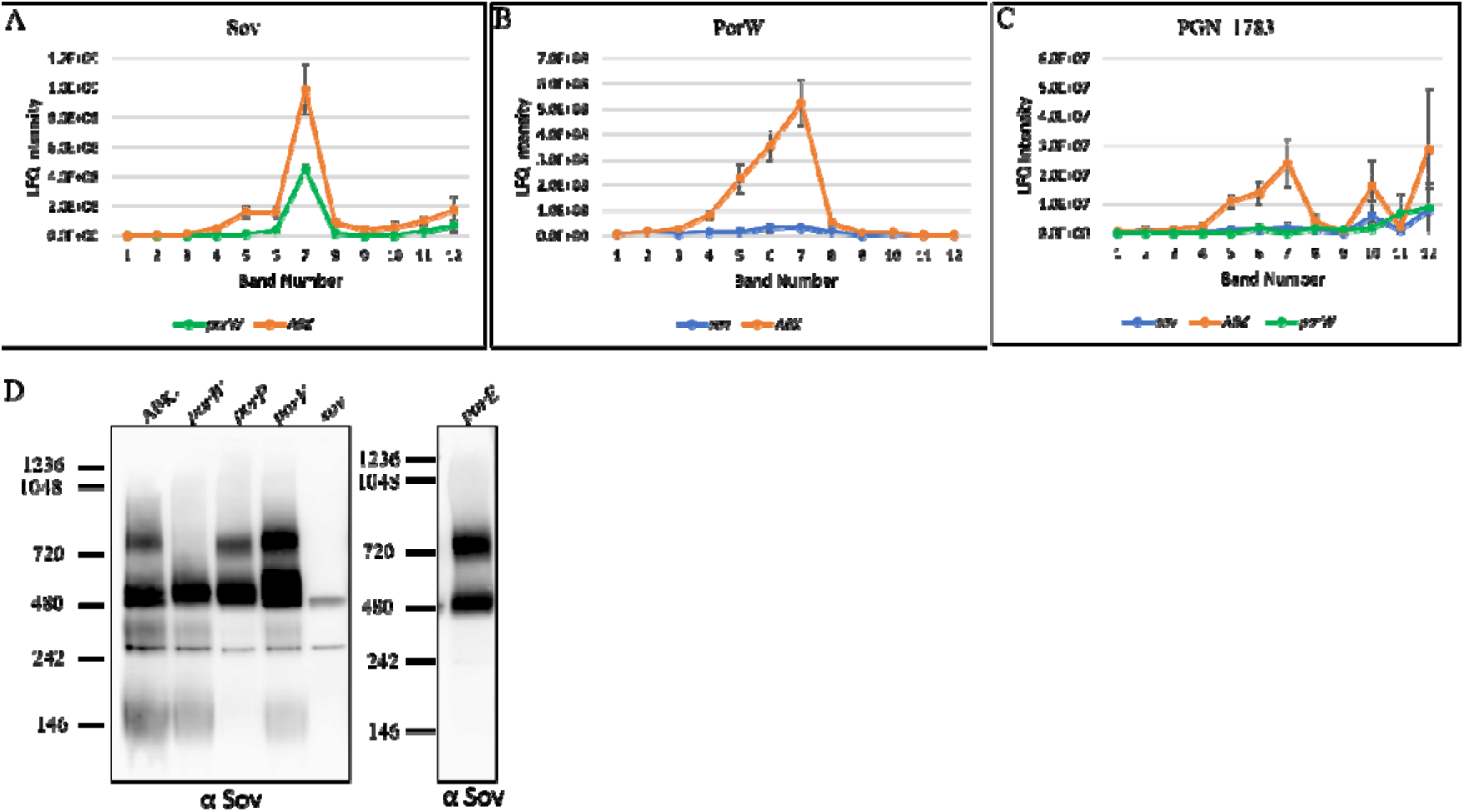
PorW is required for Sov to form the 750 kDa complex. *P. gingivalis sov* and *porW* mutants were lysed in 1% DDM and electrophoresed on a 3-12% BN-PAGE gel. The gel lanes were sliced at the same positions as in Fig 2A. The gel lanes were subjected to in-gel tryptic digestion and the tryptic fragments were analysed using LC MS/MS. The LFQ intensities of (A) Sov, (B) PorW and (C) PGN_1783 in ABK^−^, *porW* and *sov* mutants were plotted. (D) The *P. gingivalis* T9SS mutants (*porW*, *porP*, *porV*, *sov* and *porE*) and ABK^−^ were lysed in 1% DDM and electrophoresed on a BN-PAGE gel. The proteins were transferred onto a PVDF membrane and probed with Sov-specific antibodies. The *sov* mutant served as a negative control for the non-specific binding of the Sov-specific antibodies.

To obtain better resolution of the Sov complexes we performed BN-PAGE immunoblot using Sov-specific antibodies on the *P. gingivalis porW*, *porV*, *porP* and *porE* mutants (Figure 5D). These strains were selected for further analysis as they had differences in the intensities of peaks at gel bands 5 and 7 compared to the ABK^−^ strain. As anticipated, we detected two bands, one at a molecular weight of ∼500 kDa and another of ∼ 750 kDa, consistent with the BN-PAGE proteomic results. The immunoblot of *porP*, *porE* and *porV* mutants showed the same two bands as ABK^−^ (Figure 5D). In these mutant strains the signal at 500 kDa appeared to be more intense than at 750 kDa except for *porE* where it appeared to be of the same intensity consistent with the proteomics result (**Supplementary Figure 1**). Nevertheless, the band at 750 kDa was detected in all strains except for the *porW* mutant, confirming that PorW is essential for the formation of the 750 kDa Sov complex.

### Sov associates with multiple components of the T9SS

To further investigate the PPIs of Sov, we performed co-immunoprecipitation on the *P. gingivalis* ABK^−^ strain using Sov-specific antibodies. The immunoprecipitated material was quantified by mass spectrometry and MaxQuant relative to the *sov* mutant negative control. The LFQ intensity ratios of ABK^−^ relative to *sov* mutants of the top 30 proteins were plotted (Figure 6A). The highest LFQ intensity ratios were of Sov, Plug, PorW, PGN_1783, PGN_0123 and PorN (Figure 6A). PGN_0123 was recently demonstrated to be a component of the T9SS and has been named PorA (Dr. Hideharu Yukitake, personal communication). In addition to this, other T9SS components such as PorZ, PorK, PorV and PorE also had high LFQ intensity ratios (Figure 6A). The specific enrichment of many T9SS components in the co-IP suggests a large T9SS interactome involving Sov. Besides PorA several other CTD proteins such as PGN_0335 (CPG70), PGN_0659 (Hbp35), PGN_1115, PGN_1416 (PepK) and PGN_1556 (adhesin) also had high LFQ intensity ratios, consistent with their secretion through the Sov pore (Figure 6A, CTD proteins indicated with an *). To estimate the relative abundance of the proteins that co-precipitated with Sov, the iBAQ metric was used. After correction of the iBAQ values using the *sov* mutant negative control, PorV and PorA were the most abundant partners at 35% and 22% relative to the level of Sov with all other interacting components being < 4% (Figure 6B, Table 2). To determine whether the interactions changed in mutant strains, a panel of mutants comprising *porT*, *porP*, *porW* and *porV* were also subjected to Sov antibody immunoprecipitation. Similar proteins to ABK^−^ were immunoprecipitated in the *porT* and *porP* mutants, except that small amounts of the additional components PorT and PorU were found in the *porP* mutant (Table 1). For the *porW* mutant, all components identified in ABK^−^ were present except for PGN_1783, consistent with the Sov-PorW-PGN_1783 complex proposed above and suggesting that PGN_1783 is dependent on PorW for its interaction with Sov (Table 2). PorK and PorN were substantially reduced in the *porW* mutant (LFQ ratio <10) suggesting they may also depend on PorW for their interaction with Sov (Table 2). In the *porV* mutant, PorA was completely absent indicating its interaction with Sov requires PorV (Table 2). In other mutants, PorV was present at up to 1:1 stoichiometry with Sov, while the plug protein reached a maximum of 7%. In the absence of PorV, the stoichiometry of the plug rose to 140% (Table 2). PorT and PorG were also found to have high LFQ intensity ratios in the *porV* mutant suggesting they also associate directly or indirectly with Sov (Table 2). Since the proposed Sov-PorW-PGN_1783 complex is located at band 5, band 7 presumably corresponds to the Sov-PorV-PorA complex and also the Sov-PorV-Plug complex. The native MW profiles of the Plug protein (PGN_0144) confirmed its increased abundance in band 7 of the *porV* mutant sample (**Supplementary Figure 2**). Collectively, these data suggest that the Sov translocon complex is tethered to the PorK/N rings. Figure 6C shows the proposed Sov complexes of the wild-type and are discussed further below.

**Table 2:** Sov binding partners in several T9SS mutants

|  | **ABK<sup>-</sup>** |  | ***porP*** |  | ***porT*** |  | ***porW*** |  | ***porV*** |  |
| --- | --- | --- | --- | --- | --- | --- | --- | --- | --- | --- |
| **Protein Name** | **LFQ ratio** | **iBAQ %** | **LFQ ratio** | **iBAQ %** | **LFQ ratio** | **iBAQ %** | **LFQ ratio** | **iBAQ %** | **LFQ ratio** | **iBAQ %** |
| **Sov** | 4472 | 100.0 | 19121 | 100.0 | 4504 | 100.0 | 6120 | 100.0 | 283 | 100.0 |
| **PorV** | 18 | 35.3 | 264 | 77.4 | 56 | 104.4 | 64 | 93.6 | 0 | 0.0 |
| **PorA** | 98 | 22.1 | 3639 | 69.2 | 370 | 99.7 | 394 | 60.4 | 0 | 0.0 |
| **Plug** | 319 | 3.3 | 978 | 6.9 | 159 | 1.5 | 183 | 1.4 | 2305 | 142.1 |
| **PGN_1783** | 172 | 2.2 | 10 | 5.2 | 353 | 4.5 | 0 | 0.0 | 160.8 | 7.9 |
| **PorK** | 26 | 1.6 | 226 | 2.2 | 38 | 4.5 | 3 | 0.4 | 925 | 5.2 |
| **PorN** | 80 | 1.1 | 15 | 0.7 | 238 | 2.8 | 6 | 0.1 | 268 | 2.5 |
| **PorW** | 205 | 0.7 | 1220 | 2.7 | 433 | 1.3 | 0 | 0.0 | 822 | 1.9 |
| **PorZ** | 42 | 0.3 | 0 | 0.0 | 0 | 0.0 | 0 | 0.0 | 0 | 0.0 |
| **PorE** | 10 | 0.1 | 0 | 0.0 | 0 | 0.0 | 0 | 0.0 | 0 | 0.0 |
| **PorT** | 0 | 0.0 | 28 | 0.2 | 0 | 0.0 | 0 | 0.0 | 61.8 | 0.6 |
| **PorG** | 0 | 0.0 | 0 | 0.0 | 0 | 0.0 | 0 | 0.0 | 44 | 0.5 |
| **PorU** | 0 | 0.0 | 28 | 0.1 | 0 | 0.0 | 0 | 0.0 | 9.1 | 0.02 |

Note: LFQ ratios deemed highly significant (ratio $\geq 10$ ) are shaded grey.

**Figure 6:**
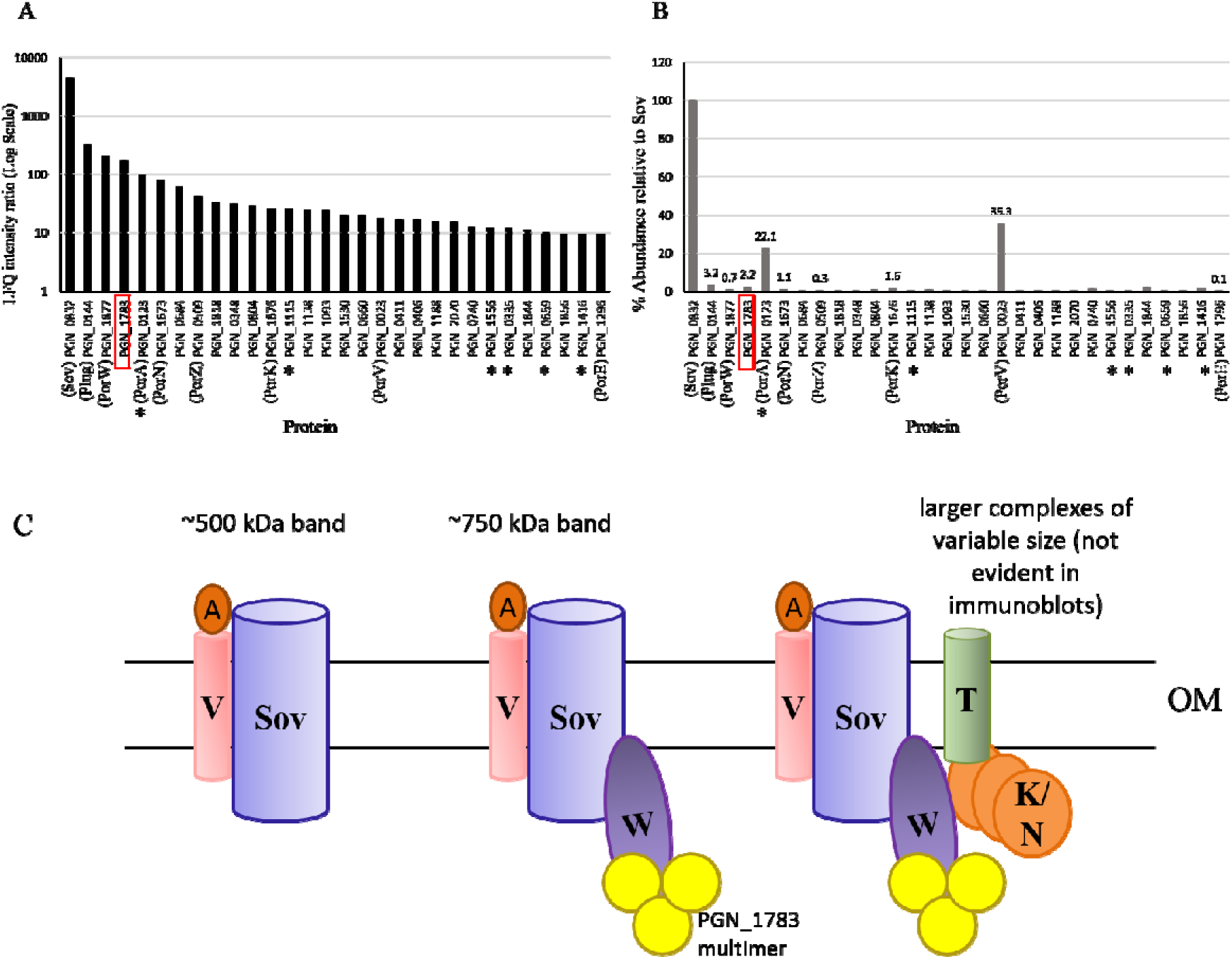
Sov interacts with multiple components of the T9SS. The *P. gingivalis* ABK^−^ strain was subjected to co-immunoprecipitation using Sov-specific antibodies. The *sov* mutant was used as a negative control. The immunoprecipitated samples were digested with trypsin and analysed by mass spectrometry and quantified using MaxQuant software. (A) The ratio of LFQ intensities (ABK^−^/sov) of the top 30 proteins was plotted. CTD proteins are marked with *. (B) For the same 30 proteins, the percentage abundance relative to Sov using iBAQ intensities was plotted. (C) Summary of the Sov proposed complexes involving Sov. Note: to show a Sov ratio in (A), the lowest LFQ intensity identified in the pull down of the *sov* mutant was used instead of a zero.

### PorP, PorE and PG1035 form a complex

To identify the potential binding partners of PorE, we expressed Strep tagged PorE (PorE^Strep^) in the *P. gingivalis* W50 *porE* mutant. This strain was black pigmented on a blood agar plate similar to WT (Figure 7A) indicating the PorE-Strep fusion was functional and supported secretion by the T9SS. Strep-tag affinity capture was performed using a cell lysate prepared from the PorE^Strep^ strain and the captured proteins were quantitated by mass spectrometry and MaxQuant relative to the WT W50 control. The LFQ intensity ratio of PorE^Strep^:WT of the top 10 proteins were plotted (Figure 7B). The LFQ intensity ratios of PorE, PorP and PG1035 were much higher than any other protein suggesting these three proteins form a complex. The iBAQ abundance of PorP and PG1035 were 1:1 whilst PorE was 10-fold more. The most plausible explanation for this is perhaps not all Strep tagged PorE proteins were able to complex with PorP and PG1035. To ensure that the increased abundance of PorP and PG1035 was due to specific pull-down and not merely due to a potential increase in the expression of these three proteins in the tagged strain, their overall abundance was quantified (**Supplementary Figure 3**). While the abundance of PorE was higher in PorE^Strep^, the abundances of PorP and PG1035 were similar in both strains, indicating that the high abundance of PorP and PG1035 in the pull down was due to association with PorE.

**Figure 7:**
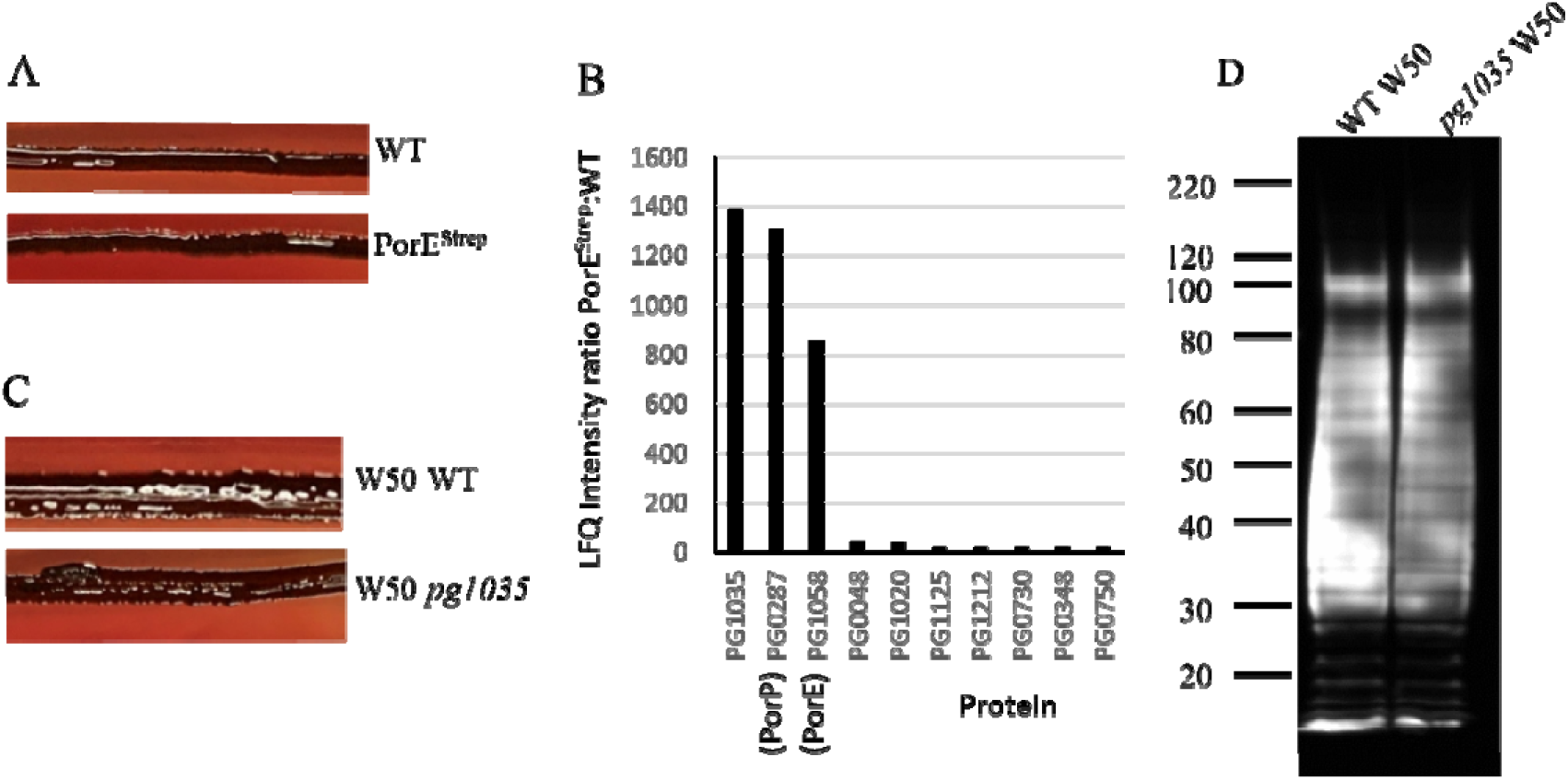
PorP, PorE and PG1035 form a complex. (A) Black pigmentation of *P. gingivalis* PorE^Strep^ strain on blood agar plate. (B) *P. gingivalis* WT and PorE^Strep^ tagged strains were subjected to affinity capture using StrepTactin sepharose beads. The affinity captured proteins were digested with trypsin and analysed by mass spectrometry and quantified using MaxQuant software. The ratio of LFQ intensities of PorE^Strep^ to WT was plotted. (C) Black pigmentation of WT and *pg1035* deletion mutant on blood agar plate (D) Proteins in the whole cell lysates of WT and *pg1035* deletion mutant were electrophoresed on an SDS-PAGE gel. The proteins were transferred onto a membrane and probed with mAb-1B5 (anti-A-LPS) antibodies.

We deleted *pg1035* in *P. gingivalis* W50 to determine if PG1035 is required for T9SS function. The *pg1035* mutant was black pigmented (Figure 7C) and an immunoblot with A-LPS antibodies showed high molecular weight A-LPS similar to WT (Figure 7D), indicating that the CTD proteins were modified with A-LPS. Thus, although PG1035 bound to PorP and PorE, it was not essential for T9SS function under the conditions tested.

To investigate which T9SS components are involved in assembly of the PorE-PorP-PG1035 complex, the native MW profiles of these three proteins in T9SS mutants was explored using both the MS data and immunoblot analysis with antibodies against PorE. The native profiles of PG1035 were vastly different between the background strains used (Figure 8A-B), indicating that the expression of PG1035 is strain-dependent. Therefore, the profiles of PorE and PorP are also presented according to their strain background (Figure 8C-F). The profiles of PorP were of poorer quality due to the low abundance (LFQ intensity) of this protein. For all mutants derived from our lab ATCC 33277 wild type strain (*porU*, *porV*, *porZ*), the three proteins tended to peak in gel band 8 (Figure 8B, D, F) which corresponded to a major band at ∼420 kDa in the native immunoblots (Figure 8G). In contrast, for mutants derived from the ATCC 33277 wild type strain obtained from Koji Nakayama (Nagasaki University, Japan), PG1035 was barely detectable, and accordingly, PorE and PorP were mainly detected at a lower MW in gel bands 9-10 (Figure 8C, E), which in the native immunoblots corresponded to the major band at ∼260 kDa likely to represent a PorE-PorP complex (Figure 8G). This was confirmed by direct comparison of the W50 *pg1035* mutant with other W50 strains (Figure 8H). In the strains expressing PG1035 (W50 ABK^−^, W50 *porU*), the ∼420 kDa band was produced corresponding to the PorE-PorP-PG1035 complex, while in the absence of PG1035, only the ∼260 kDa complex comprising PorE-PorP was produced. The near absence of PG1035 in the strains obtained from Koji Nakayama was also confirmed in the ATCC 33277 wild type strain.

**Figure 8:**
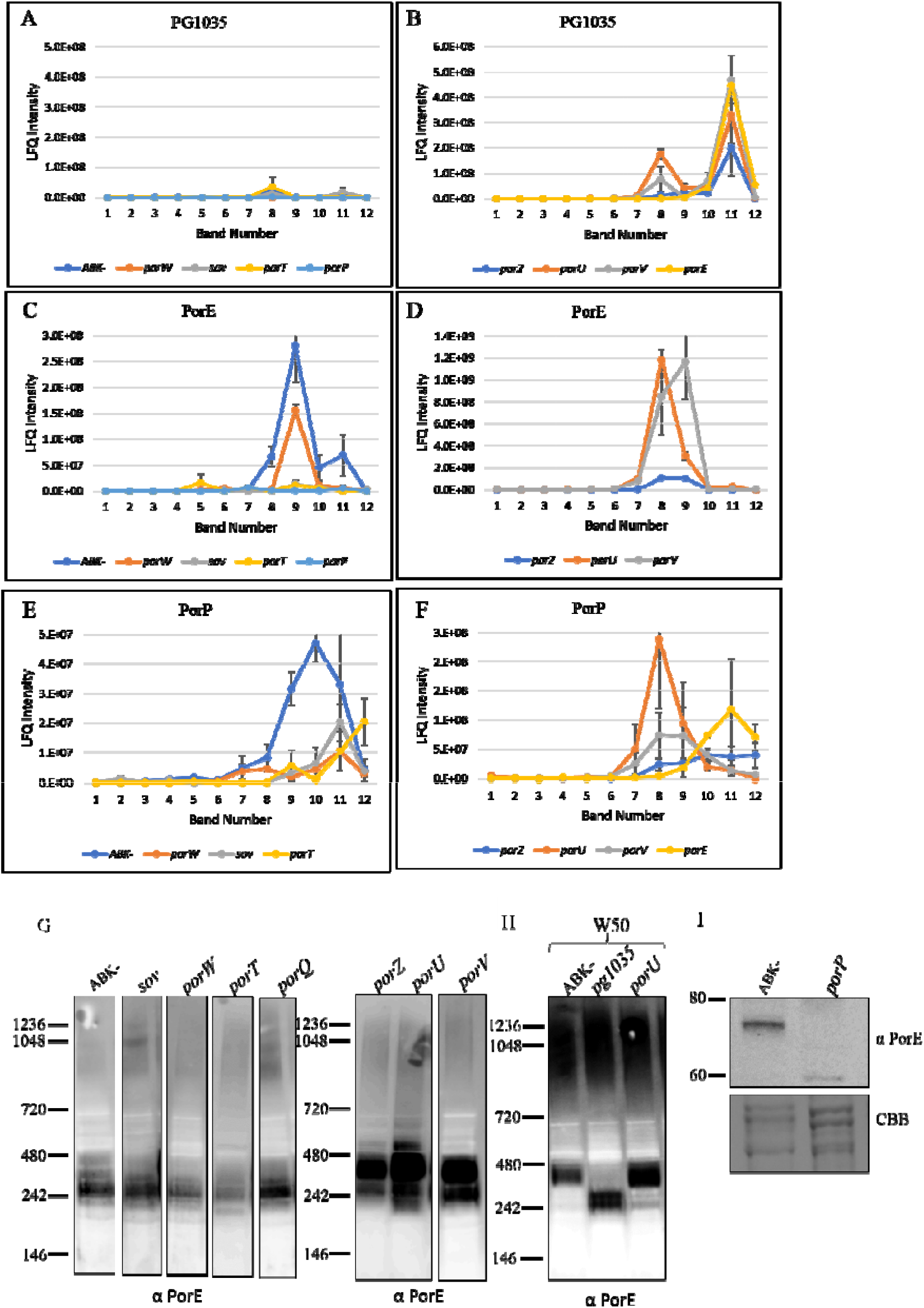
PorP stabilises PorE and strain-specific native migration profiles of PorP, PorE and PG1035. *P. gingivalis* T9SS mutants (*porT*, *porP*, *porE*, *porW*, *sov*, *porZ*, *porU* and *porV*) were lysed in 1% DDM and electrophoresed on a 3-12% BN-PAGE gel. The gel lanes were sliced as in Fig 2A. The tryptic fragments were analysed by mass spectrometry. The LFQ intensities of PG1035(A-B), PorE (C-D) and PorP (E-F) were plotted for each T9SS *P. gingivalis* mutant and ABK^−^. The strains plotted on the left (A, C, E) are the ATCC 33277 obtained from the Nakayama laboratory while the strains plotted on the right (B, D, F) are the ATCC 33277 from our laboratory. (G) Cell lysates from the selected strains were electrophoresed on a BN-PAGE gel and the proteins were transferred onto a PVDF membrane and were probed with PorE antibodies. (H) Whole cell lysates from ABK^−^, *pg1035* and *porU* mutants from *P. gingivalis* W50 background were electrophoresed on a BN-PAGE gel and proteins were transferred onto a PVDF membrane and probed with anti-PorE antibodies. (I) Whole cell lysates from *P. gingivalis* ABK^−^ and *porP* mutant were electrophoresed on an SDS-PAGE gel. The proteins were transferred onto a nitrocellulose membrane and probed with anti-PorE antibodies. Coomassie blue (CBB) stained gel shows the relative loading amount.

PorE was detected in all the mutants in both strain backgrounds except for the *porP* mutant suggesting that PorE may be unstable in this mutant (Figure 8C). Denaturing immunoblot analysis on the *porP* mutant using antibodies specific for PorE confirmed this suggestion. In ABK^−,^ PorE was detected at its expected MW of ∼78 kDa, while in the *porP* mutant only a faint band was detected at ∼60 kDa (Figure 8I). This suggests that in the absence of PorP, PorE is unstable and is degraded. In the W50 *porE* mutant, PorP and PG1035 were both found at a lower native size peaking at band 11 (Figure 8B, F), possibly representing a PorP-PG1035 complex. The native size of PorF was unaffected in both *porP* and *porE* mutants (**Supplementary Figure 4**) indicating that it is not complexed with either PorP or PorE, consistent with the absence of PorF in the PorE^Strep^ pull down (Figure 7B).

### The *F. johnsoniae* motility protein SprB binds to the PorP homologue SprF

*P. gingivalis* has only one protein that has a type B CTD, PG1035. Secretion of PG1035 has not been studied but as shown here, it appears to form a complex with PorP and PorE. *F. johnsoniae* type B CTD protein SprB is co-expressed with the PorP orthologue, SprF, which is essential for SprB secretion and cell-surface localization ^15^. What SprB interacts with on the cell surface and how SprB interacts with the motility machinery is not known. Since the *P. gingivalis* type B CTD protein PG1035 is likely bound to PorP on *P. gingivalis* cell-surface, we reasoned that SprB may be bound to the PorP orthologue SprF, on the *F. johnsoniae* cell surface. To test this hypothesis, we performed co-immunoprecipitation on a *F. johnsoniae* strain co-expressing (i) SprF and (ii) the CTD of SprB fused to sfGFP (sfGFP-CTD_SprB (368 aa)_) ^15^. The sfGFP in this strain was previously shown to be rapidly propelled along the cell surface demonstrating its successful secretion by the T9SS and association with the motility machinery ^15^. The material immunoprecipitated by anti-GFP was quantitated by mass spectrometry and MaxQuant relative to a WT control. The LFQ intensity ratios and iBAQ abundances of sfGFP-CTD_SprB (368 aa)_ relative to WT were plotted for the top 30 proteins (Figure 9A and B). The highest LFQ intensity ratios were of SprF and sfGFP-CTD_SprB (368 aa)_ suggesting SprF and SprB interact directly through the SprB CTD. The iBAQ abundances were similar suggesting a 1:1 stoichiometry between these two proteins. Other known components of the gliding/T9SS machinery such as SprA, GldJ, Fjoh_4997 (PPI), SprE, GldN and GldK, as well as the putative new component, Fjoh_3466 (homologue of PGN_1783, which as shown above appears to interact with Sov and PorW) were also specifically pulled down, but at much lower abundance (Figure 9A and 9B). Figure 9C summarises the apparent interactions of SprB associated proteins. Together, these results suggest that SprB may be held on the surface and brought into contact with the rest of the motility machinery via a direct interaction with the PorP orthologue, SprF. The interaction of type B CTD protein PG1035 with PorP and of type B CTD protein SprB with PorP-like protein SprF, suggests that other type B CTD proteins may also interact with their appropriate PorP-like proteins on the cell surface.

**Figure 9:**
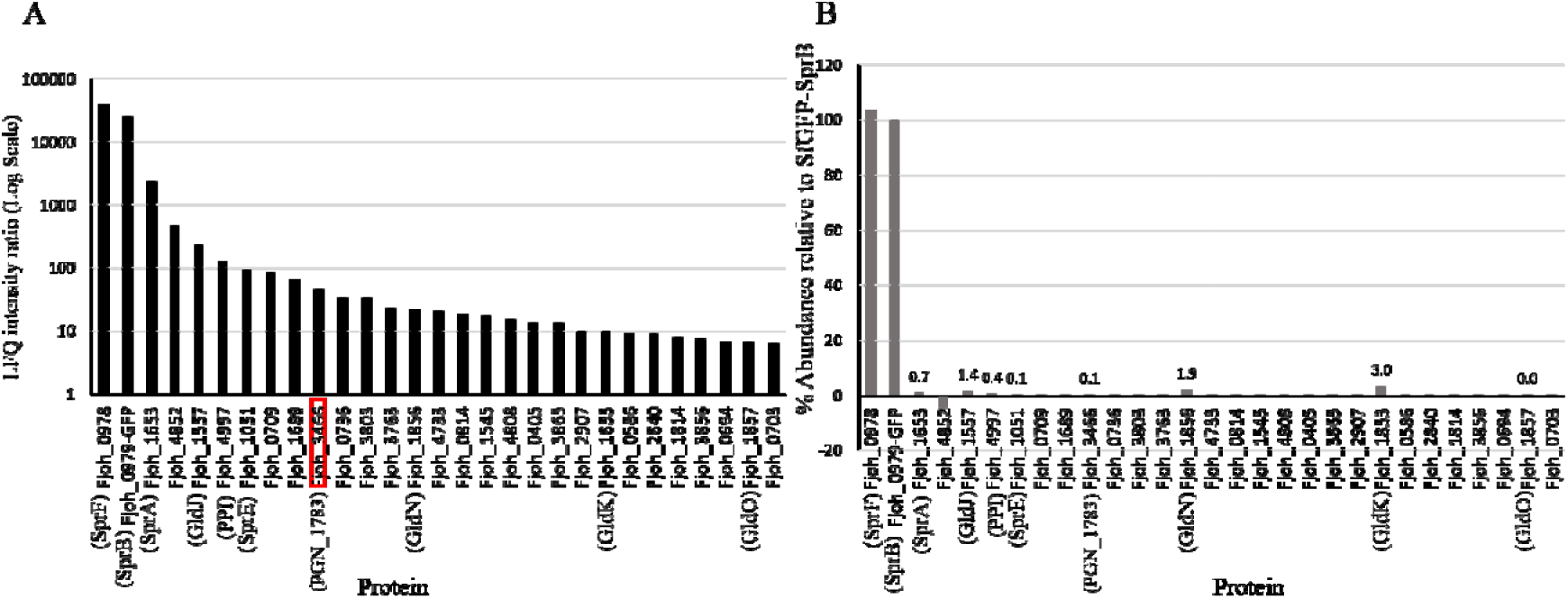
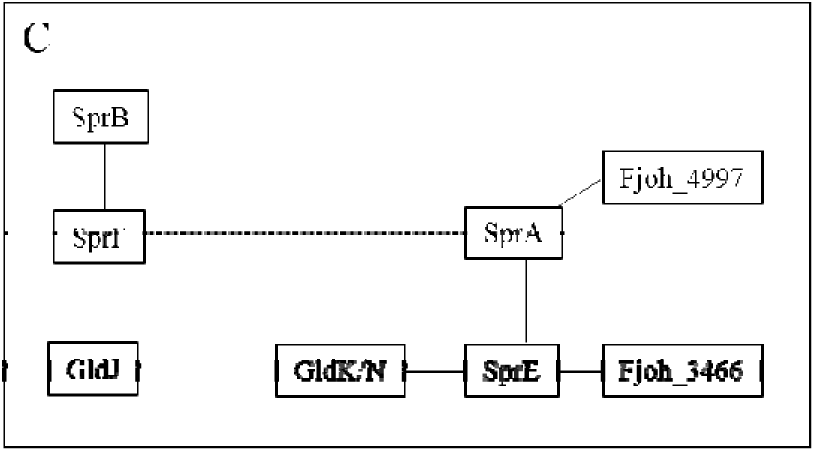
SprB and SprF form a complex. *F. johnsoniae* wildtype and sfGFP-CTD_SprB (368 aa)_ strains were subjected to co-immunoprecipitation with GFP agarose beads. The immunoprecipitated samples were digested with trypsin and analysed by mass spectrometry and quantified using MaxQuant software. (A) The ratio of LFQ intensities of sfGFP-CTD_SprB (368 aa)_ to WT and (B) % abundance based on iBAQ intensities were plotted. (C) Summary of the protein interactions.

## Discussion

In this study, we present the first *in situ* cryo-ET images of the *P. gingivalis* T9SS. The cryotomograms show that the T9SS forms large rings of 50 nm diameter in the bacterial periplasm immediately adjacent to the outer membrane. Our previous single particle reconstruction of purified PorK-PorN also revealed 50 nm diameter rings formed from the assembly of 32-36 monomers of each protein ^25^. It has been questioned whether the rings may have been an artifact of purification without PorM or PorL rather than a native structure ^27^. Our cryotomograms of *P. gingivalis* cells with fully functional T9SS reveal that the large PorK-PorN rings are a native structure. While there are other densities near PorK-PorN that appear faintly above the noise, we were unable to align or resolve other components of the T9SS. This is likely due to heterogeneity in the particles, either in the stage of assembly, in the functional state, or in the inherent disorder of other periplasmic components.

We therefore turned to proteomic approaches to explore the organisation of the PorK/N rings with respect to the other T9SS components. We show that these rings are part of the *P. gingivalis* core translocation machinery comprising at least 10 components namely PorK, PorN, PorG, PorT, PGN_1783, PorW, Sov, PorV, PorA and Plug. This core machinery may interact transiently with the PorL/M and PorP/E/PG1035 complexes, while the attachment complex appears to be detached.

The core translocation machinery was built up in the following way. The PorK/N/G complex in *P. gingivalis* ^25^ and the SprA/PorV and SprA/Plug complexes in *F. johnsoniae* ^24^ were already demonstrated. Our co-IP data using anti-Sov showed that Sov (SprA orthologue) also interacted with PorV and Plug in *P. gingivalis.* In addition, it was found that PorA was part of this complex at a similar stoichiometry as PorV, and that its inclusion in the complex was dependent on PorV. The BN-PAGE proteomics data together with the co-IP data indicated that Sov was also bound to the PorW lipoprotein and PGN_1783. Furthermore, Sov and PorW are present at a similar abundance in *P. gingivalis*, suggesting these two proteins are present at a similar stoichiometry ^47^. This complex could be observed in immunoblots at ∼750 kDa. PorK/N together with PGN_1783 were pulled down in WT and all mutant strains tested except for *porW*, suggesting that PorW binds directly to Sov, while PorK/N and PGN_1783 are bound to PorW. PorK and PorN form large ring complexes with a BN-PAGE MW of > 4 MDa ^25^, therefore the 750 kDa band cannot include intact PorK/N complexes. Since PorK and PorN were strongly detected in BN bands 1-5, it is likely that the ring proteins had dissociated to some extent, resulting in complexes of different size. The 750 kDa Sov complex is therefore likely to be a distinct complex that includes PorW and PGN_1783 while additional associations with PorK and PorN may be smeared in the background over a wide MW range with different sized ring fragments (Figure 6C).

The presence of PorT in the complex was also supported by both BN-PAGE and co-IP experiments. Since its BN profile corresponded best with PorK/N, it is likely to be bound directly to the rings in low stoichiometry, similar to PorG ^25^ (Figure 10A). The core translocation machinery is likely to be similar in *F. johnsoniae*, as in the sfGFP-CTD_SprB(368_ aa) pull down, many of the same components were detected, including SprA, SprE, GldK/N and Fjoh_3466 which are orthologues of Sov, PorW, PorK/N and PGN_1783 respectively.

**Figure 10:**
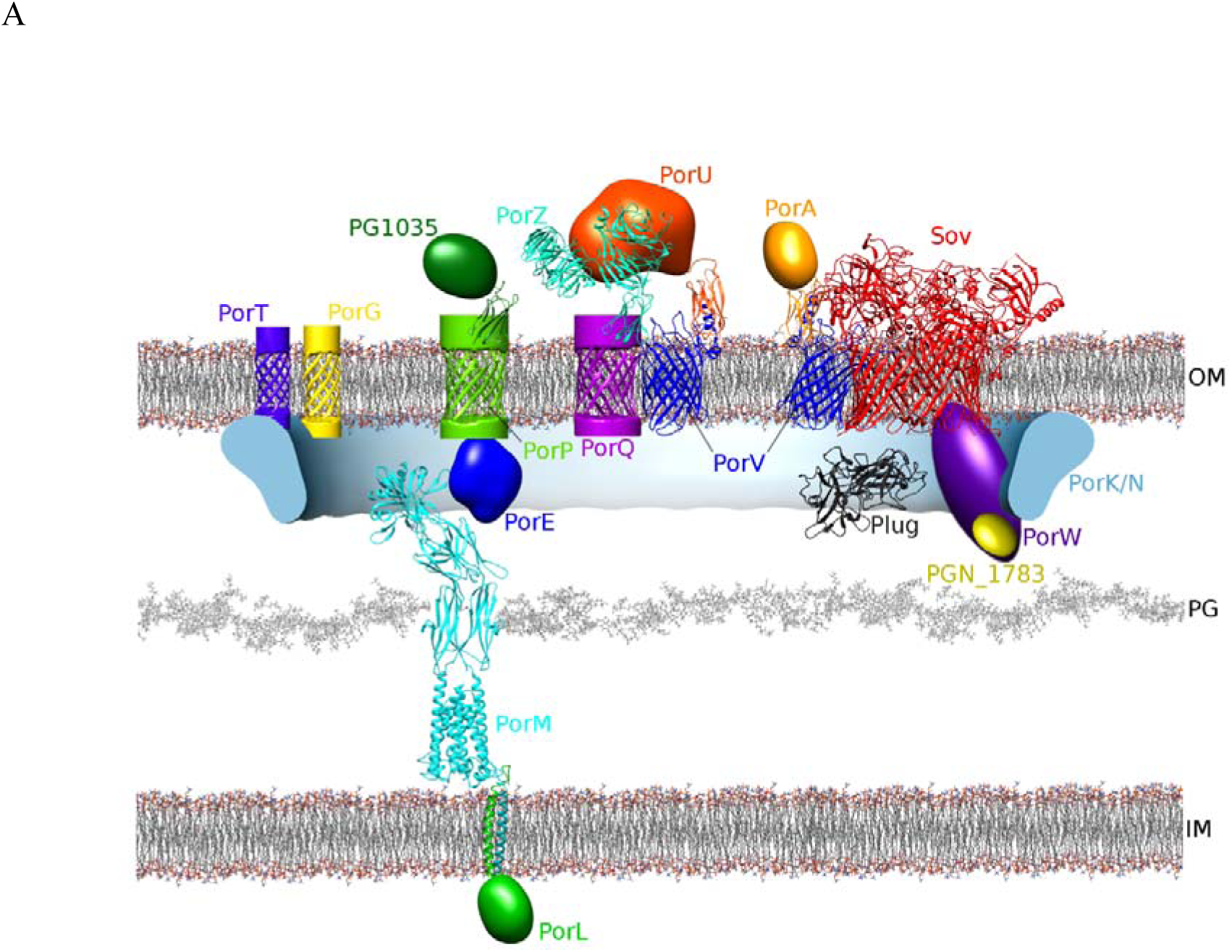
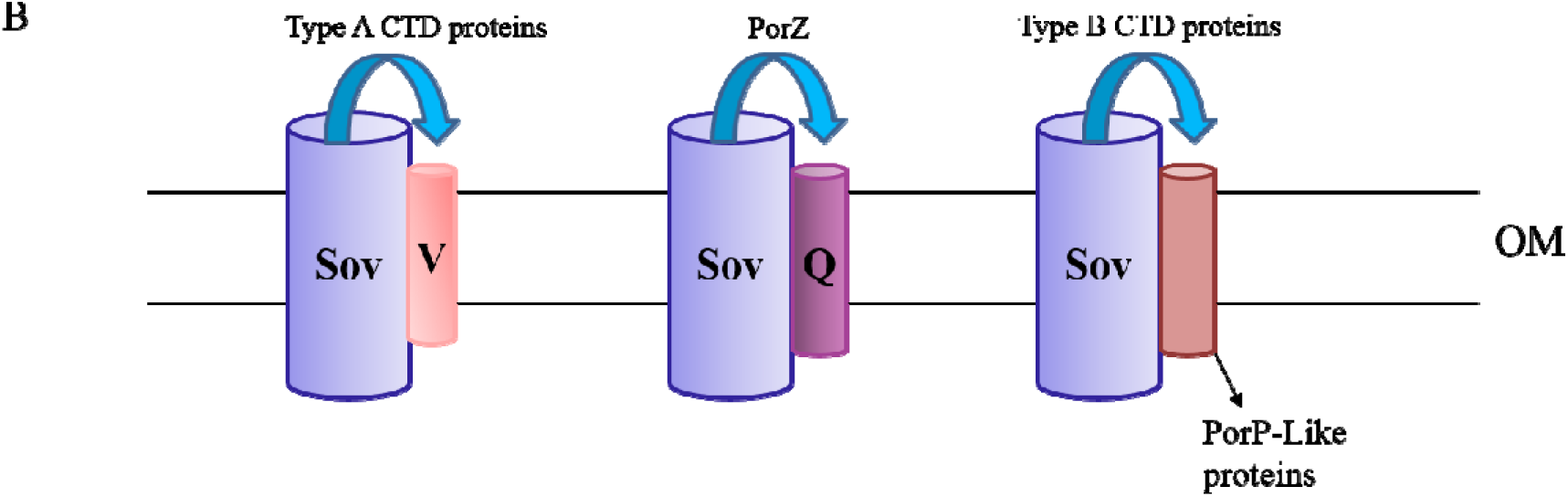
Proposed organisation of the T9SS components within the PorK/N rings. (A) Proposed structural organization of the T9SS. The Sov translocon interacts directly with PorV, Plug and PorW. PorA and PGN_1783 bind to the translocation machinery via PorV and PorW, respectively. PorW/PGN_1783 connects the Sov translocon to the PorK/N rings that also interact with PorG and PorT. PorP, PorE and PG1035 form a complex that may interact transiently with the translocation machinery as well as with the PorL and PorM complex in the inner membrane. The attachment complex consisting of PorU, PorV, PorQ and PorZ appears to be independent and does not associate directly with the other T9SS complexes. We propose all these sub-complexes to be localised inside the PorK/N rings so that PorV can collect the cargo proteins from Sov and shuttle them to the attachment complex for anchorage to the cell surface. The schematic diagram does not show the stoichiometry of each component. PorQ and PorP are predicted 14-stranded beta barrels similar to PorV, while PorT and PorG are predicted 8-stranded beta barrels. The PDB entries (www.rcsb.org) for the structures shown are: Sov/PorV (6H3I), Plug (6H3J), PorZ (5M11), PorM (combination of 6EY5 and 6EY0), PorQ and PorP barrels used PorV (6H3I) and PorT and PorG barrels used OmpA (1BXW). Molecular graphics were produced with UCSF Chimera^51^. (B) Schematic diagram showing proposed association of PorV, PorQ and PorP-like proteins with the Sov translocon to collect their respective cargo proteins.

In a previous screening analysis, a *P. gingivalis* PGN_1783 deletion mutant was black pigmented, indicating a functional T9SS ^3^. This mutant should be re-examined in more detail to determine if PGN_1783 has a detectable role in the T9SS, especially since some other T9SS components, including the Plug and PG1035, appear to have non-essential roles under laboratory conditions. It was unexpected to find such a close association between Sov-PorV-PorA, which suggests a specialised role for PorA in the function of the T9SS. PorV interacts with type A CTDs on the surface ^28^, and thus may interact with PorA via its type A CTD. As the Sov-PorV-PorA complex was also found in the T9SS mutants, this suggests that PorA can still translocate through Sov to reach PorV on the surface in the absence of a fully functional T9SS. Alternatively, the presence of the Sov-PorV-PorA complex in the T9SS mutants may indicate that PorA interacts with the Sov-PorV complex in the periplasm, but this option seems less likely. PorA has recently been proposed to be a functional component of the T9SS since the *porA* mutant was non-pigmented (Dr. Hideharu Yukitake, personal communication). While PorA was present in the pull downs at high LFQ ratios and also at high abundance (iBAQ%) in all strains except the *porV* mutant, other type A CTD proteins were either absent or present at much lower levels in all strains.

Our BN-PAGE proteomics study also revealed that the surface attachment complex proteins PorQ and PorZ form a sub-complex in the absence of PorU or PorV. Glew et al., identified a complex of PorZ with the OM beta barrel protein PorQ in the PorUC690A catalytic mutant providing a mechanism for PorZ to be anchored to the cell surface ^28^. Previously, Lasica et al., showed that PorZ is on the cell surface in the absence of PorU ^20^. Therefore, it is likely that PorZ is on the surface bound to PorQ in the *porV* mutant as well. Together, it appears that PorZ can translocate through the T9SS secretion channel in the absence of PorU or PorV. PorU and PorZ both have conserved CTD domains and are known to be T9SS substrates as well as components. However, unlike other CTD proteins, the CTD signals of PorU and PorZ are not cleaved ^17, 20^. Kulkarni et al., have shown that there are at least two types of CTD signal, type A and type B ^14^. *F. johnsoniae* type A CTD proteins are dependent on PorV for secretion whereas many type B CTD proteins are not ^48 15^. PorZ appears to have a CTD that is neither type A or type B. Since PorV forms a complex with Sov ^24^ and binds to newly secreted CTD proteins before shuttling them to the attachment complex ^28^, we suggest that both PorQ and PorV target the same Sov binding site and recruit their respective substrates accordingly.

A novel complex was identified consisting of PorP, PorE and PG1035. PorP is an OM beta barrel while PorE is an OM associated lipoprotein localised in the periplasm with a predicted peptidoglycan-binding domain ^3, 19^. PG1035 is the only *P. gingivalis* protein with a type B CTD. Given that all proteins containing either type A or type B CTD are secreted and anchored onto the cell surface or released in the culture fluid, it is likely that PG1035 is translocated across the OM and attached to the cell surface via PorP. The anchorage to PorP is likely to be through the CTD of PG1035 as its CTD was observed to be intact in the pull-down.

The CTD of PG1035 is similar to the CTD of *F. johnsoniae* SprB, and those of 11 other *F. johnsoniae* type B CTD proteins. In contrast to *P. gingivalis*, which encodes a single PorP, *F. johnsoniae* encodes 11 PorP-like proteins, and these are thought to be specific for secretion of their cognate type B CTD proteins ^15, 48^. SprB, for example, requires its cognate PorP-like protein SprF for secretion and attachment on the cell surface. SprB is a highly repetitive 669 kDa protein that moves rapidly along the cell surface, resulting in gliding motility. How this protein interacts with the secretion and motility machine(s) is not known. We have shown that sfGFP-CTD_SprB (368 aa)_ and SprF form a complex and the interaction is via the CTD of SprB. The interaction could be either at the periplasmic side of the OM or on the cell surface. Given that GFP was observed on the cell surface ^15^ it is likely the interaction occurs there. Together, this suggests that SprB is anchored to the cell surface through SprF. The motility protein GldJ was also specifically pulled down with SprB, but at a lower abundance than SprF. GldJ is an OM lipoprotein that is required for propulsion of SprB on the cell surface. It appears to localise in a helical manner, and it was proposed to be associated with the track on which SprB is propelled ^49 50^. SprF may link SprB, directly or indirectly, to this track. Based on our data and the gene arrangement in *F. johnsoniae* where most SprF-like proteins are located immediately downstream of genes encoding type B CTDs ^15^ we postulate that other SprF-like proteins may also anchor their respective type B CTD proteins to the cell surface.

SprB was also found associated with SprA, Fjoh_4997, Fjoh_3466, SprE, GldN and GldK (Figure 9). These interactions may have been captured during the translocation of SprB through the SprA channel. Previously, SprA was shown to interact with the peptidyl-prolyl *cis-trans* isomerase, Fjoh_4997 ^24^ which is consistent with our findings. SprA is known to associate with either PorV or the Plug ^24^, but in the SprB pull-down neither of these proteins were observed (Figure 9C). This may not be surprising since SprB secretion is independent of PorV and instead requires SprF ^48^. PorV and SprF may bind to the same site on SprA to collect their respective cargo proteins. When SprB is being secreted, SprF, rather than PorV, would be complexed with SprA (Figure 10B).

The demonstration that PorK/N forms 50 nm rings *in situ* raises interesting questions regarding their function. The ring is continuous and packed against the OM in a way that might prevent the diffusion of trans-membrane proteins across it and out of the ring. Also, it appears that the T9SS includes spatially separate OM complexes and hence there may be a functional need for them to stay in close proximity. An important function of the rings therefore may be to keep these separate protein components close together within the outer membrane. Cargo proteins are first directed to the Sov translocon, and then secreted and bound to PorV on the surface. We propose that this occurs inside the rings. As the PorV shuttle transports the cargo to the attachment complexes, these attachment complexes should also be located within the rings. Once cargo is transferred from the attachment complex to an A-LPS anchor located in the outer leaflet of the OM, the cargo protein would then be free to pass beyond the PorK/N rings to form the EDSL. Since the expression of PorK and PorN is essential for T9SS function, the barrier may be essential for the correct localisation and assembly of the other components. The need for such a ring in the T9SS, but not in other secretion systems may be associated with the presence of the attachment complexes and shuttle system which need to be in close proximity to the secretion pore.

In summary, we show an *in situ* structure of the T9SS and have identified a comprehensive interactome of *P. gingivalis* T9SS components (Figure 10A). While secreting substrates with a type A CTD signal, we propose that the Sov translocon at any one time, is either bound to PorV/PorA or to the plug and is localised within the PorK/N rings and connected to them via PorW and PGN_1783. The small 8-stranded OM β-barrels PorG and PorT are associated with the PorK/N rings. Together, these 10 components are proposed to comprise the core translocation machinery (Figure 10A). The attachment complex comprising PorU-V-Q-Z appears to be independent of the other complexes (Figure 10A). PorV, PorQ and the PorP-like proteins are all 14-stranded OM β-barrels of the FadL family that independently bind to cargo proteins with a specific type of CTD signal. We therefore propose that each one is able to collect its respective substrate protein(s) directly from the Sov translocon (Figure 10B). We also propose that the role of PorK/N rings is to form a barrier for all the OM T9SS components to contain them as an OM island allowing harmonized secretion and cell surface attachment of the T9SS substrates. Together, the results provide the first *in situ* structure of the T9SS and important insights regarding its organisation and architecture.

## Acknowledgements

We acknowledge the use of the Mass Spectrometry and Proteomics Facility at the Bio21 Institute, The University of Melbourne, Australia. This work was supported by the Australian National Health and Medical Research Council grant ID 1123866, the Australian Government Department of Industry, Innovation and Science Grant ID 20080108 and the Australian Dental Research Foundation (ADRF) Grant ID 349-2018. This work was also supported by NIH GRANT A127401 to GJJ and NSF GRANT MCB-1516990 to MJM. Cryo-EM work was done at the Beckman Institute Resource Center for Transmission Electron Microscopy at Caltech.

