## Supplementary Tables 1 and 3, Supplementary Figs 1-4 for "*In situ* structure and organisation of the type IX secretion system"

**Supplementary Information**

**Supplementary Table 1: List of *P. gingivalis* strains used in this study**

| ***P. gingivalis* strain** | **Source** |
| --- | --- |
| 33277 ABK- | Koji Nakayama laboratory |
| 33277 *porW* | Koji Nakayama laboratory |
| 33277 *porP* | Koji Nakayama laboratory |
| 33277 *porQ* | Koji Nakayama laboratory |
| 33277 *porT* | Koji Nakayama laboratory |
| 33277 *sov* | Koji Nakayama laboratory |
| 33277 *porK* | Koji Nakayama laboratory |
| 33277 *porU* | Laboratory collection |
| 33277 *porV* | Laboratory collection |
| 33277 *porZ* | Laboratory collection |
| W50 *porE* | Laboratory collection |
| W50 ABK- | Laboratory collection |
| W50 *porU* | Laboratory collection |
| W50 *pg1035* | Laboratory collection |
| W50ABK^-^*WbaP | Laboratory collection |

**Supplementary Table 3: LFQ intensities (the highest intensity in each mutant have been assigned 100%) of PorG in *porP*, *porE*, *porU* and *porV* *P.gingivalis* mutants; PorG showed a similar migration profile to PorK and PorN**

|  | ***porE*** | ***porP*** | ***porU*** | ***porV*** |
| --- | --- | --- | --- | --- |
| **Band 1** | 100 | 89 | 96 | 79 |
| **Band 2** | 35 | 62 | 11 | 55 |
| **Band 3** | 13 | 9 | 0 | 16 |
| **Band 4** | 45 | 100 | 56 | 44 |
| **Band 5** | 33 | 31 | 18 | 45 |
| **Band 6** | 13 | 6 | 0 | 18 |
| **Band 7** | 2 | 0 | 0 | 0 |
| **Band 8** | 3 | 13 | 100 | 8 |
| **Band 9** | 0 | 22 | 0 | 16 |
| **Band 10** | 37 | 96 | 24 | 63 |
| **Band 11** | 32 | 69 | 13 | 100 |
| **Band 12** | 26 | 88 | 0 | 79 |

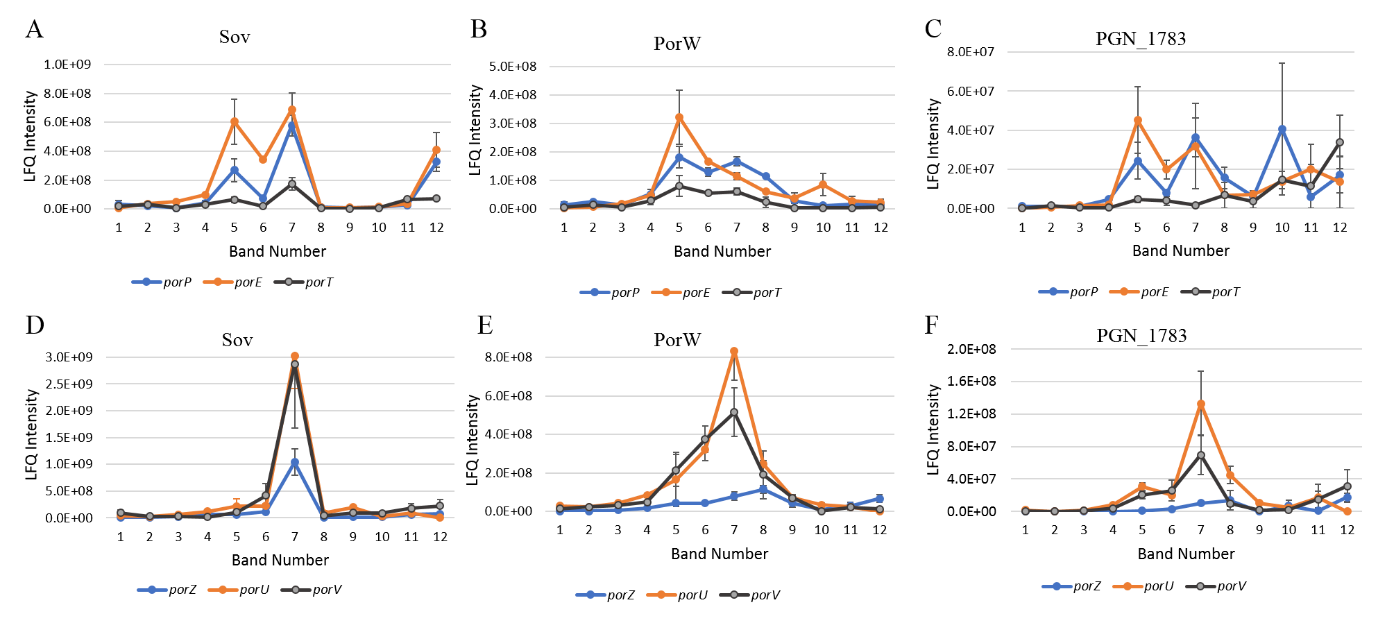

**Supplementary Figure 1: Correlation of Sov-PorW-PGN_1783 native profiles**

*P. gingivalis* T9SS mutants (*porT*, *porP*, *porE*, *porZ*, *porU* and *porV*) were lysed in 1% DDM and electrophoresed on a 3-12% BN-PAGE gel. The gel lanes were sliced at the same positions as in **Fig 1A**. The gel lanes were subjected to in-gel tryptic digestion and the tryptic fragments were analysed by mass spectrometry. The LFQ intensities of Sov, PorW and PGN_1783 in each T9SS mutants were plotted. Sov plots were grouped based on whether a Sov peak was predominant at Band 5 (A) or Band 7 (D). PorW (B and E) and PGN_1783 (C and F) groups were based on Sov grouping.

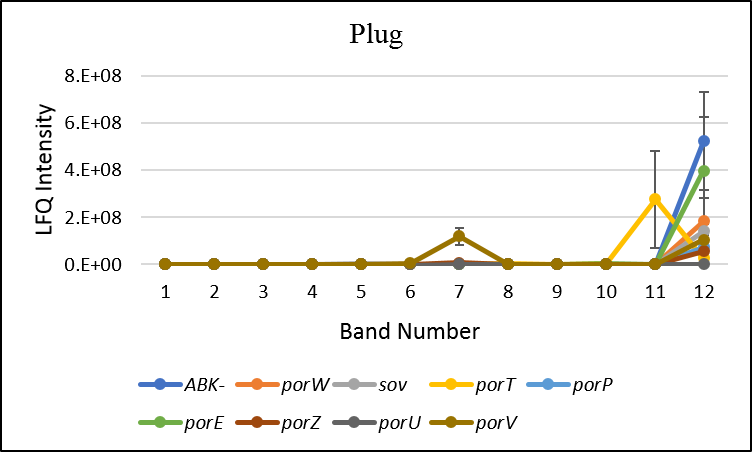

**Supplementary Figure 2: Plug protein at band 7 in *porV* mutant**

*P. gingivalis* T9SS mutants (*porT*, *porP*, *porE*, *sov, porW*, *porZ*, *porU* and *porV*) were lysed in 1% DDM and electrophoresed on a 3-12% BN-PAGE gel. The gel lanes were sliced as in Fig 1A. The gel lanes were subjected to in-gel tryptic digestion and the tryptic fragments were analysed by mass spectrometry. The LFQ intensities of the Plug protein in each T9SS mutants was plotted

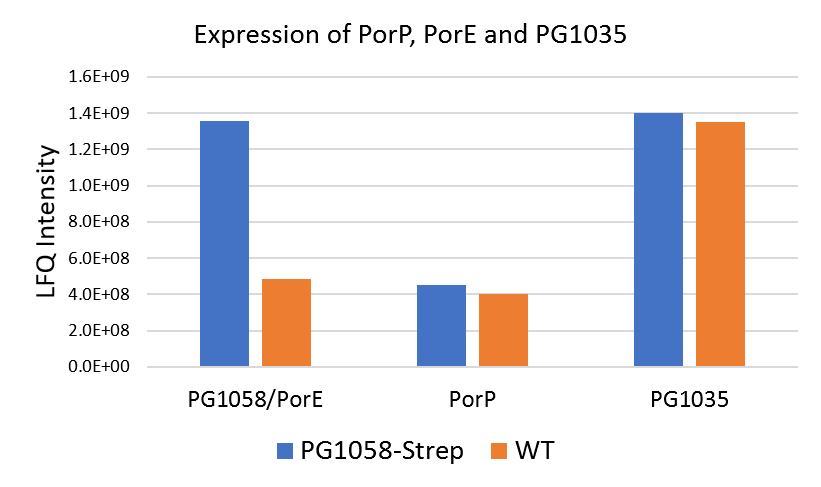

**Supplementary Figure 3: Overall abundance of PorE, PorP and PG1035 in WT and PorE^Strep^ *P. gingivalis* W50**

Cell lysates from WT and PorE^Strep^ *P. gingivalis* cells were electrophoresed on SDS-PAGE for a short time. The sample in each lane was excised as a single band and subjected to in-gel tryptic digestion. The tryptic fragments were analysed by mass spectrometry. The LFQ intensity of PorE, PorP and PG1035 in WT and PorE^Strep^ are plotted.

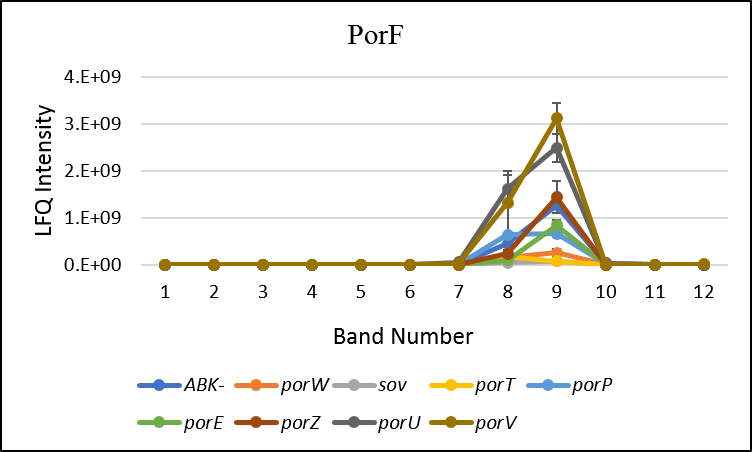

**Supplementary Figure 4: Native migration profile of PorF in T9SS mutants**

*P. gingivalis* T9SS mutants (*porT*, *porQ*, *porP*, *porE*, *sov, porW*, *porZ*, *porU* and *porV*) were lysed in 1% DDM and electrophoresed on a 3-12% BN-PAGE gel. The gel lanes were sliced as in Fig 1A. The gel lanes were subjected to in-gel tryptic digestion and the tryptic fragments were analysed using MaxQuant software. The LFQ intensities of PorF protein in each T9SS mutants were plotted
